# The compounding costs of being female in academia: Individual-based modelling of career progression and interventions

**DOI:** 10.64898/2026.04.29.721520

**Authors:** Liam Gibson, Ann Brower, Lindsey Te Ata o Tū MacDonald, Alex James

**Affiliations:** School of Earth and Environment, University of Canterbury, Christchurch, New Zealand; School of Mathematics and Statistics, University of Canterbury, Christchurch, New Zealand; Bioprotection Aotearoa Centre of Research Excellence, New Zealand; School of Politics and Political Science, University of Canterbury, Christchurch, New Zealand

**Keywords:** gender equity, individual-based modelling, management practices

## Abstract

We develop a novel individual-based population dynamics model of academic career progression, using 15 years of data from over 1,000 academics from one university. Our model improves on previous models, which, by homogenising career progression, may underestimate the costs of being female.

We find multiple effects that compound to slow career progression for women. Women are hired at lower ranks than men, then face the sticky floor problem of getting stuck at the bottom for longer. Further, individuals in STEM fields are promoted more quickly; this disproportionately affects women who are more prevalent in non-STEM fields.

Our model reveals age is more complicated than others have found with ODE-based models. Women are older when hired, and promotions favour the young; hence age costs women more. Finally, the probability of attrition rises with years spent at the same rank, regardless of gender. Since women are promoted slower, they experience higher attrition rates.

We also deploy our model to test possible interventions. We find just hiring more women will not work. A more nuanced set of interventions is required. Gender parity will only be achieved at the highest ranks if hiring rates are jointly equalised across gender, academic rank, and discipline.

## 1. Introduction

Academia has a gender problem. Countless research papers and reporting projects have found that women are underrepresented at the highest ranks of academia [1-7]. Gendered rank disparities lead to institutional gender pay gaps, even when women and men are paid the same for the same role [8]. Motivated by both representation and pay equity, closing these gender gaps is a goal for many universities [9, 10].

Previous studies have used age to explain the gender problem [11-13]. The argument for the age cause is twofold – demographics compounded by gendered research patterns. Demographic inertia observes that men tend to be older [14] because previous generations of academia had more men [3, 11, 15]; the theory proposes that the demographics and age-induced gender gaps will sort themselves out as we hire more early career women [6, 7, 16, 17]. Further, older academics tend to publish more than younger academics [13, 18, 19]; ergo, men tend to publish more than women [11, 13, 18, 20] (and reap the associated promotion/pay benefits). Delving into various forms of age reveals it is not so cut and dry. Many researchers have found that the number of years since the start of an academic career [20, 21] and the number of years spent at a given academic rank [15] are more important than biological age. Studies which do find biological age to be important in academic publication often uncover a complex relationship with productivity [19, 22]. For example, there is evidence to suggest that academic productivity may slow down after a certain age threshold [19, 23]; older academics may take a more supervisory role [22]; in some disciplines, older academics may prefer legacy-style publication types, such as books [21]; women and men may have different relationships between age and productivity [24].

Importantly for us, the main objection to the ‘age explanation’ is that there are more senior women in academia than ever before, yet gender gaps in academia have not yet closed (nor are they projected to without institutional intervention [4-6, 25, 26]). This suggests that there are more factors at play than biological age.

We start with the premise that there are three key drivers of faculty composition: hiring, promotion, and attrition. Recent studies highlight the importance of gender balance in academic hiring [2, 4, 7, 27]. LaBerge et al. [27] find that hiring impacts faculty composition more than attrition in over 90% of academic disciplines, despite the same data set showing that women have higher attrition rates compared to men [28]. James & Brower [4] show that disparities in hiring rate and rank (i.e., the number of women hired and their academic rank upon hire) affect gender representation gaps more than promotion and attrition. Thomas et al. [7] show that gender parity for STEM faculty is achieved only via hiring more tenured women, ensuring they are retained and promoted at rates equal to men. Just as there is more to gender gaps than age, we see more than hiring as possible interventions.

Barrett-Walker et al. [1] find that higher attrition rates among women lead to shorter careers. To earn the same amount over a lifetime, women would have to be promoted to higher ranks earlier than men. However, Brower & James [2] showed that women have a lower probability of promotion to high-ranked positions during their career compared to men with similar research scores. Using a simple model of cumulative pay, Brower & James [2] estimate an average lifetime earnings gap of NZ$400,000 between women and men.

To compare hiring, promotion, and attrition (and their interactions), we present a case study using a single institution, the University of Canterbury (UC). UC is a large public research university in Aotearoa New Zealand, with women comprising about 38% of the total academic staff in 2020. UC uses a rank-based salary scale, similar to the UK and Australia. There are four main academic ranks (Lecturer, Senior Lecturer, Associate Professor, and Professor) and individuals must apply for promotion to move to the next rank. In 2020, at the highest academic rank (Professor), women were outnumbered by 2.6 to 1, meaning about 73% of Professors were men.

We explore the roles of age, discipline, and rank in academic progression, gender representation, and gender pay gaps. We use those results to model various interventions. Simply hiring more women is not the quick fix of long-term gender gaps, nor will hiring practices alone fix the problem of gender gaps in pay and representation. More nuanced combinations of interventions are required to narrow gaps in gender representation and pay. This is at least partly due to the role of discipline, simplified here to STEM and non-STEM. The most effective combination of interventions would remove any explicit gender bias in promotion and attrition, equalise gender balance in hiring, minimise the gender gap in rank at hiring, and narrow the gender representation gap in STEM.

## 2. Data

We use 8,943 payroll records from the University of Canterbury (UC) from between 2005-2020. This data set contains the academic rank (Lecturer, Senior Lecturer, Associate Professor, Professor), pay step within the rank, birth year, gender, and broad discipline (by Faculty of Science, Arts, Education, Engineering, Business, and Law) from *N* = 1,107 anonymised staff. Each staff member is assigned a unique ID number allowing individuals to be tracked through their career. Academics outside these ranks (e.g. postdoctoral researchers, tutors, Assistant Lecturers, etc) are not included in the analysis. Each rank has between 4 and 8 pay steps within it and, at lower ranks, individuals move automatically up these steps each year (see Supplementary Material for rank and step details). All staff have the opportunity to apply for promotion each year to progress to the next rank, regardless of which pay step they are on. Double rank promotions (e.g. from Lecturer to Associate Professor) are theoretically possible, but none were recorded during this time. Disciplines within the Faculties of Engineering and Science were categorised as STEM. Disciplines from the Faculties of Education, Business, Law and Arts, were classified as non-STEM. A more detailed data description and summary are in Supplementary Materials.

While useful, the University of Canterbury data set has limitations. There is no method to disaggregate sex and gender, nor is data collected on non-binary academics. Transgender individuals were assumed to have their preferred gender, e.g., trans men were included as men. Individuals with missing gender data and gender queer academics are both assigned ‘unknown’ gender in the data set. These academics (about 1% of the total data set) are excluded from our analysis. This is a large limitation, so we acknowledge here that LGBTQ academics (in STEM fields) tend to have fewer career opportunities and resources, and more thoughts and plans to leave academia [29]. This results in higher attrition rates for junior LGBTQ academics, on average [30]. Further, given the limited ethnicity information in the University of Canterbury data set (one priority ethnicity per academic and many missing entries), we do not consider ethnicity here. Barrett-Walker et al. have shown that marginalised ethnicity groups have significantly higher attrition rates compared to European academics [1].

The number of women hired per year has increased during the 15 years of the data set by 21.0%. Accordingly, there is a 35.6% decrease in men’s hiring. In an average year, the Lecturer rank is comprised of 47.4% men and 52.6% women. Women make up 37.5% of Senior Lecturers, 29.4% of Associate Professors, and 20.5% of Professors. At the Lecturer and Senior Lecturer ranks, 45.6% and 28.1% of new hires are women, respectively. At the Associate Professor rank, this increases to 44.2% but reduces back to 15.7% for Professors. Supplementary Materials, Table S1, Figure S1, and Figure S2 present these data in more detail.

Between years 2005 and 2020, the average male Lecturer has increased from 39.4 years old to 42.3 years old, and the average female Lecturer age has increased from 39.3 years old to 44.0 years old. Female Senior Lecturers were, on average 48.4 years old in 2005, increasing to 49.2 years old in 2020, conversely the average age of male Senior Lecturers decreased from 48.5 to 47.3 over the same period. This is similar at the Associate Professor rank, where women’s average age increased from 48.7 to 51.5 years old, and men’s average age decreased from 53.6 to 51.4. Finally, the Professor rank has the largest change in age over the data period, with the average woman’s age increasing from 51.5 to 56.1 and men decreasing from 57.6 to 56.2. See Supplementary Material (Table S1 and Figure S3) for more details.

We find subtle differences in hiring age between women and men at every academic rank. This is clearest at the Lecturer rank, where a substantial number of hires have been made. Lecturer hiring age distributions are unimodal with a long tail extending into the high age range. This indicates that most Lecturers are hired between ages 30 and 40 but can be hired well into their 50s. Women tend to make up a larger proportion of these older hires compared to men, with a higher mean age of 39 (compared to men’s 36.8) and a higher standard deviation of 8.5 (compared to men’s 6.9) (Figure S4). Other ranks have less well-defined structures. Overall, women tend to be hired about 2 years older than men. Finally, women are more likely to be hired as Lecturers in comparison to men. This is true across all years of the data set and in both STEM and non-STEM disciplines. See Supplementary Materials for details.

Academic salaries are dictated by the Tertiary Education Union (TEU) Collective Employment Agreement [8] which maps academic rank and step to annual salary in NZD. At lower ranks, individuals automatically move up a single step within their rank each year and apply for promotion when they have reached the top step for that rank. To avoid inflation-related issues all ranks and steps were assigned salaries using the 2022 collective agreement scale included in Supplementary Materials. In addition to the collective agreement salary, some academics receive additional salary either through private consulting work or as an individual pay agreement. These payments were not available in our data and are not included.

## 3. Statistical models

We use logistic generalised linear models to estimate the relationship between individual demographics, such as age, gender and discipline with academic promotion and attrition. We define promotion as an individual moving from one rank to the next and attrition as an individual leaving the data set. Our key independent variables are gender (*IsMale*), years since birth (*Age*), academic rank (*Rank*), years spent at that rank (*YearsAtRank*), discipline (*IsSTEM*) and being over 65 years old, i.e. retirement eligibility (*Retirement*). After a promotion an individual’s *RankAge* is reset to zero.

Using AIC, *k*-fold cross validation, and focusing on significant interactions (see Supplementary Materials for full model selection details), we use the generalised linear models (given in Wilkinson notation),

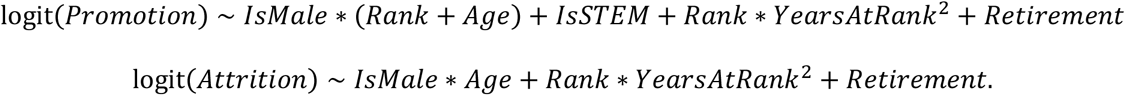

Generalised linear mixed models were used to account for individual-level random effects and found the same results. Previous research has shown that early career men may receive more mentoring and support than early career women [31, 32] which is seen here in the significance of the interaction between gender and age, and gender and rank. STEM disciplines, particularly engineering, are dominant at UC and have more external funds available to them. This is a possible cause of the positive and significant *IsSTEM* term in the promotion equation. The interaction between rank and years-at-rank-squared is understandable given that promotions are unlikely to happen in very quick succession as individuals must build up a new portfolio to get the next promotion. But conversely getting “stuck” at a rank for many years is detrimental to promotion chances too, possibly indicating the existence of a personal career ceiling.

Table 1 shows the full model output including coefficients and *p*-values. We find that men have significantly higher promotion rates than women, although this advantage decreases with age. Age is significantly negatively correlated with promotion and positively correlated with attrition. Promotion and attrition rates are significantly impacted by academic rank, and this varies with gender. Men are advantaged at the Lecturer rank but slowly lose their advantage higher up the academic hierarchy. Academic staff in STEM disciplines have significantly higher promotion odds, regardless of gender, age, rank, and all other predictors.

**Table 1:** Generalised linear models and coefficient values. Logistic regression models of promotion and attrition fit to UC data between 2005-2020. Bolded values are significant with *p* < 0.05.

| *Coefficient* | *Description* | Model |  |
| --- | --- | --- | --- |
|  |  | *Promotion* | *Attrition* |
| Male | Gender indicator | **1.30\*\*** | 0.08 |
| Age | Biological age | **-0.04\*\*\*** | **0.03\*\*** |
| SL | Senior Lecturer | **-1.17\*\*** | 0.61 |
| AP | Associate Professor | 0.26 | -0.06 |
| P | Professor | - | -0.01 |
| YearsAtRank | Years at rank | **0.81\*\*\*** | **0.36\*\*\*** |
| YearsAtRank<sup>2</sup> | Years at rank squared | **-0.06\*\*\*** | **-0.02\*\*** |
| Retirement | Retirement indicator | **-0.78\*** | **1.24\*\*\*** |
| STEM | Discipline indicator | **0.80\*\*\*** | - |
| Male:SL | Interaction between gender and rank | -0.37 | - |
| Male:AP |  | **-0.91\*\*\*** | - |
| Male:P |  | - | - |
| Male:Age | Gender/age interaction | **-0.02\*** | $-4.38 \times 10^{-3}$ |
| SL:YearsAtRank | Interaction between rank and years at rank | -0.11 | **-0.27\*\*** |
| AP:YearsAtRank |  | -0.16 | -0.21 |
| P:YearsAtRank |  | - | -0.19 |
| SL:YearsAtRank<sup>2</sup> | Interaction between rank and years at rank squared | **0.03\*\*** | **0.02\*** |
| AP:YearsAtRank<sup>2</sup> |  | 0.02 | 0.01 |
| P:YearsAtRank<sup>2</sup> |  | - | 0.01 |
| (Intercept) | - | **-2.30\*\*\*** | **-5.05\*\*\*** |

Note:
\* $p < 0.05$ ; \*\* $p < 0.01$ ; \*\*\* $p < 0.001$

## 4. Individual-based model of representation

To explore the effects of different interventions, we propose an agent- or individual-based model (IBM). This model applies the statistical model results to a set of individuals and looks at population level effects over many years.

We consider a population of *N* individuals at year *y = 0*, where each individual *i* = 1, 2, …, *N*has a designated gender, discipline (STEM or non-STEM), rank, and age. Every individual is independently predicted to leave, be promoted to the next rank, or continue at their current rank, based on the probabilities predicted by our statistical models. Promotion and attrition are independent processes. Each year, we update academic demographics, i.e., age is incremented by 1, years at rank is incremented by 1 (if no promotion occurs), and retirement eligibility is recalculated using biological age. Promoted academics’ ranks are incremented and their *YearsAtRank* variable reset to zero. Leaving academics (and academics aged over 75) are removed from the simulated population. If leaving and promotion occur in a single year, academics are removed due to attrition, but their final recorded rank is incremented. To maintain a constant population size *N*, we assume that the *N*_*A*_ individuals designated to leave are replaced by *N*_*A*_new individuals. New individuals are selected by randomly sampling *N*_*A*_IDs (with replacement) from the UC data set, and using each individual’s first year as the initial condition.

### Model output – promotion and attrition

To explore differences in promotion and attrition, we calculate the probability of ultimate promotion from Lecturer (L) to Senior Lecturer (SL), i.e., the probability that an individual hired as a Lecturer is promoted at least once during their career. To highlight demographic differences between women and men, we perform this calculation for STEM and non-STEM disciplines, and for 30-year-old and 40-year-old, newly hired Lecturers (Figure 1). Younger men in STEM fields have the highest probability of ultimately moving to Senior Lecturer (over 90%), conversely older women in non-STEM fields have the lowest chance (less than 55%).

**Figure 1:**
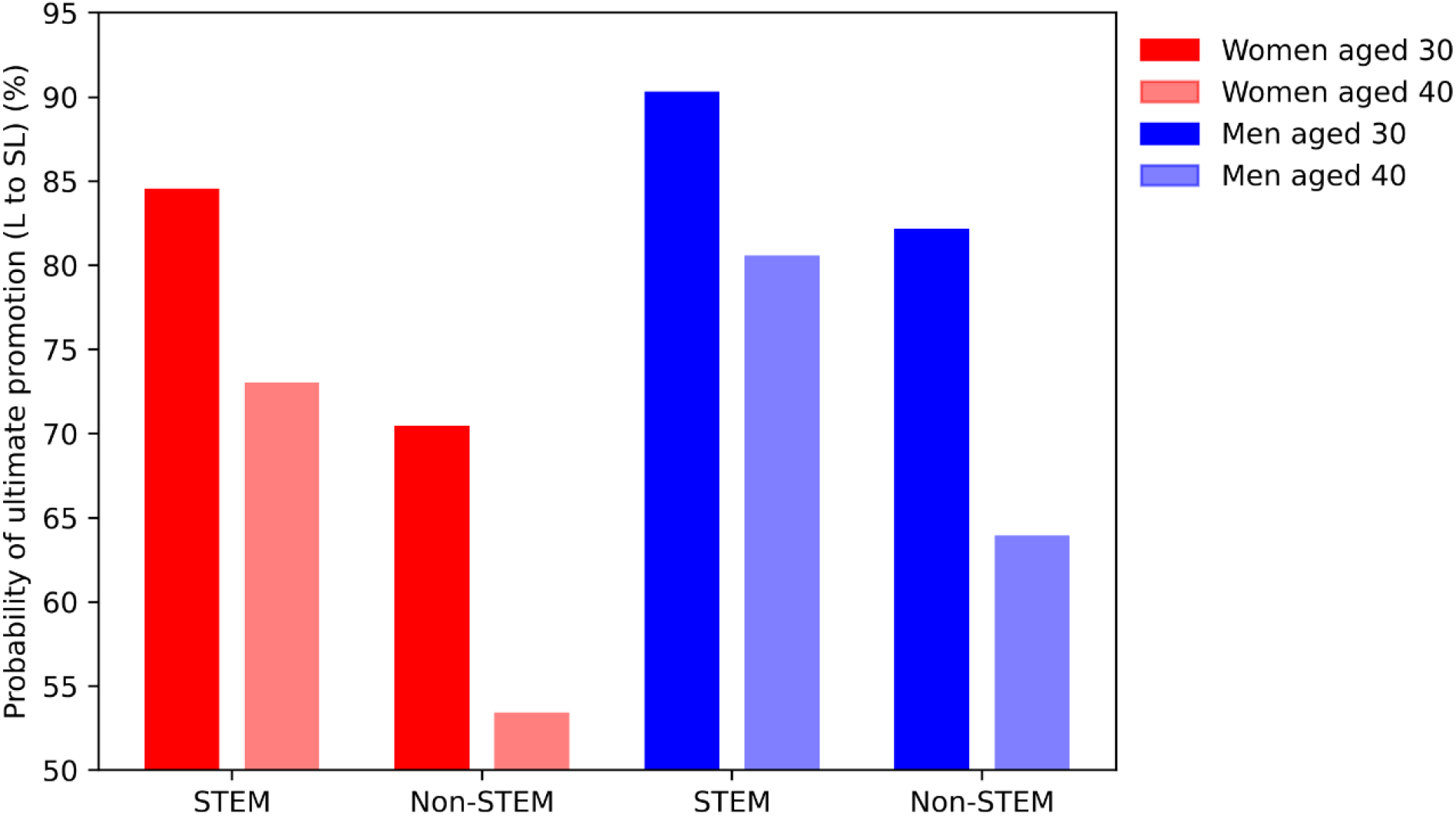
Compared to men, women are less likely to leave the Lecturer rank during their careers. Probability of ultimate promotion from Lecturer to Senior Lecturer calculated using output from generalised linear models (see Supplementary Materials for details). Bars – probabilities; red – female Lecturers aged 30; light red – female Lecturers aged 40; blue – male Lecturers aged 30; light blue – male Lecturers aged 40.

The majority of female Lecturers (61.3%) are hired in non-STEM subjects; their average hiring age is 40.5 years old. This gives them a 52.6% probability of ultimate promotion, i.e. 47.4% never reach the Senior Lecturer rank and either leave or stay at the Lecturer level until retirement. The remaining 38.7% of female Lecturers are hired in STEM subjects where only 22.6% of them never reach Senior Lecturer. Conversely, the majority of male Lecturers (61.1%) are hired in STEM subjects with an average hiring age of 35.6. Only 14.3% of these men will *not* reach Senior Lecturer. Of the remaining 38.9% of men in non-STEM, hired at an average age of 38.5, 32.9% will never reach Senior Lecturer. In addition to being more likely to get stuck at Lecturer, women are more likely to be hired at the Lecturer rank than men. Overall 70.1% of women are hired as Lecturers, in comparison to only 52.6% of men. In STEM fields this is more prominent where 80.0% of women are hired at Lecturer compared to 54.1% of men.

### Model output – validation with data

Next, we initialise the model using the UC data set. To reduce noise, records from 2005-2007 are bootstrap resampled such that the initial number of academics equals the number at UC in 2005. Figure 2 shows the averaged results of 1,000 simulations for 35 years. For the higher ranks, the model shows an excellent fit, as measured by Pearson’s *R*-squared, with the data. At lower ranks, the fit is less good, as data are driven more by annual hiring decisions than the overall system parameters. The model uses hiring data averaged over the full 15-year period, when disruptions such as COVID-19 and the Canterbury earthquakes gave large annual variance in hiring numbers. Moreover, lower ranks show weaker change over time, with the change in Lecturers failing to reach statistical significance (*p* > 0.05; time coefficient in linear regression). With no relationship between population and time, the *R*-squared metric becomes poorly defined. Although the full data set is not publicly available due to privacy concerns, Table S1 gives a summary of the initial condition and the hiring data that can be used for replication (together with Figure S5).

**Figure 2:**
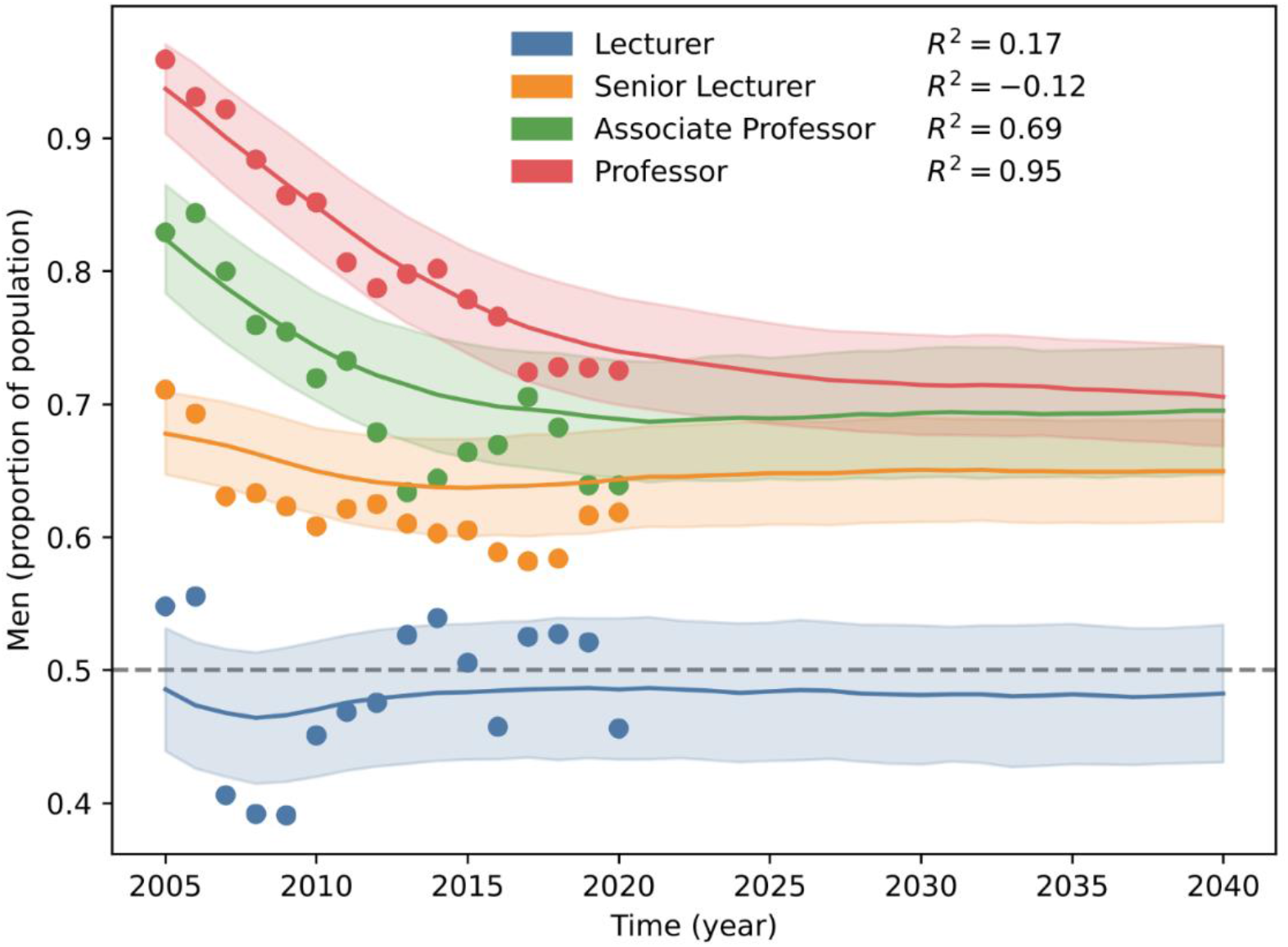
The model gives a very good fit at the higher ranks to the data. Proportion of men at each academic rank, Lecturer (L), Senior Lecturer (SL), Associate Professor (AP), and Professor (P). Initial condition is a bootstrap resampled UC data set between years 2005-2007. Dots – UC data; dashed line – gender parity; solid line – model output; blue – Lecturer rank; orange – Senior Lecturer rank; green – Associate Professor rank; red – Professor rank; shaded region – standard deviation.

### Model output – long term predictions

Finally, we initialise the model with *N* = 1,000academic IDs sampled, with replacement, from the UC data set and choose a random year in their career to create our initial condition.

The system stabilises after approximately 50 years, i.e. the longest time an individual will stay in the system and reaches a stochastic steady-state after approximately 200 years (4 full academic generations). To ignore this transient behaviour, we discard the first 500 years of the simulation results and run the simulation for an additional 2,000 years. Figure 3 shows a summary of the long-term results for the STEM, non-STEM, and overall groups. The model predicts that, based on patterns of hiring, promotion and attrition from the last 20 years, women will continue to be underrepresented at the highest academic ranks (Figure 3). On average, women are expected to comprise 52.4% of Lecturers, 35.1% of Senior Lecturers, 31.4% of Associate Professors, and 30.5% of Professors. These representation gaps are starker in STEM fields, where women make up 39.8%, 26.6%, 19.9% and 23.3% of the ranks from Lecturer to Professor, respectively.

**Figure 3:**
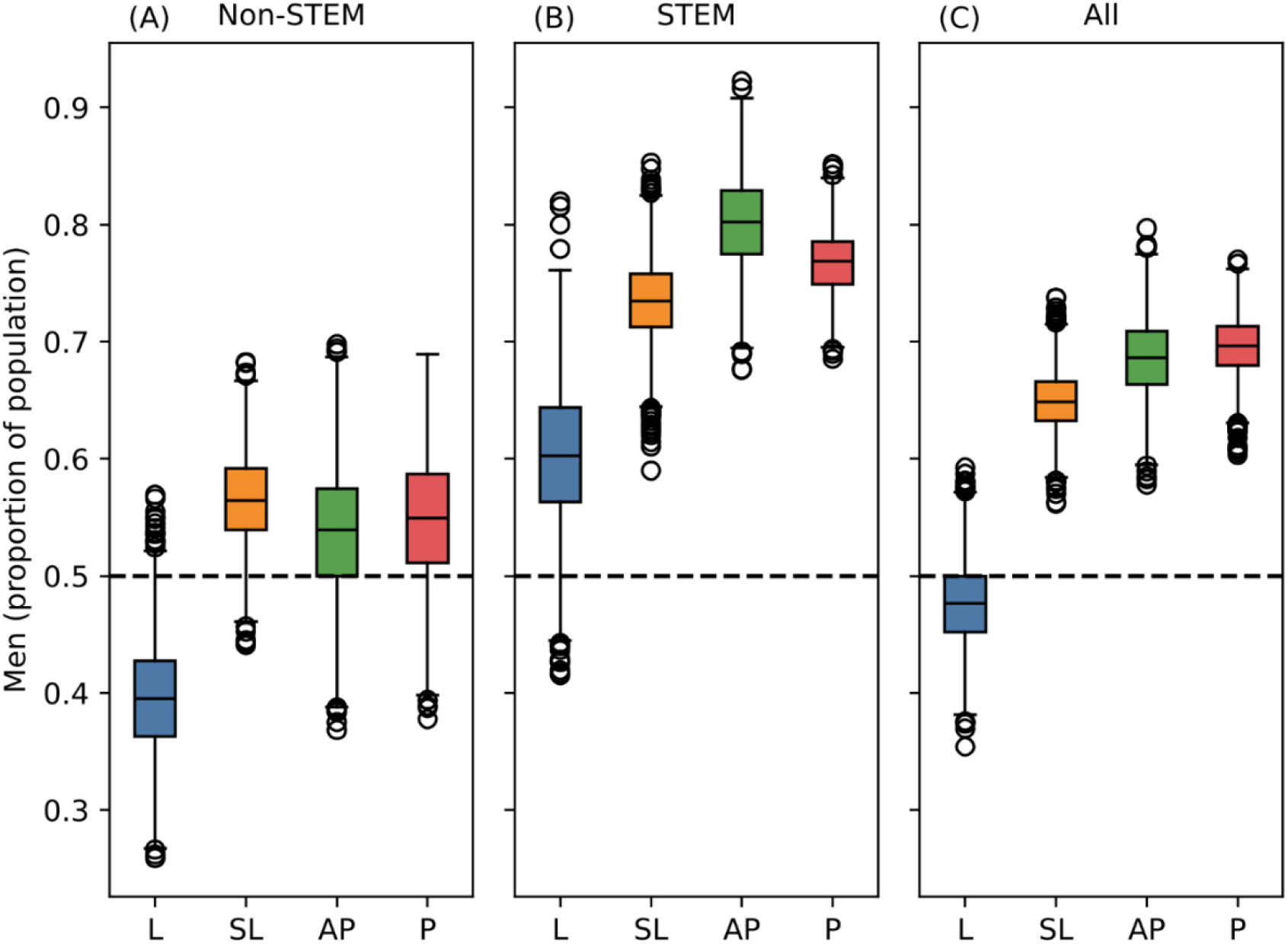
Based on current promotion, attrition, and hiring, individual-based modelling predicts that women will continue to be underrepresented in the professoriate, even after accounting for other demographics. Boxplots (median, IQR, 90^th^ percentile, outliers) showing the gender composition within each rank quantify women’s representation within non-STEM (A) and STEM (B) disciplines, and overall (C). Blue - Lecturer rank; orange - Senior Lecturer rank; green - Associate Professor rank; red - Professor rank; dashed line – gender parity.

Across all disciplines, women’s representation decreases monotonically with rank (Figure 3). However, in STEM fields, there are fewer female Associate Professors than full Professors, and in non-STEM fields, there are more female Associate Professors than Senior Lecturers. These non-linear proportion rankings are driven by the rank/gender interaction in the promotion GLM. Specifically, the decrease in men’s relative promotion advantage between Senior Lecturer and Associate Professor (Table 1). Since promotion rates in STEM fields are higher than non-STEM fields, regardless of gender and rank, STEM faculty are more likely to reach Associate Professor, where they experience this relative disadvantage.

## 5. Modelling institutional interventions

To further explore differences in hiring and other gender equity policies, we model a range of institutional interventions.

**Equal promotion** shuffles academics’ genders at random before promotion probabilities are generated, then reassigns them to their original gender after promotion decisions are made. This does not give full equality in promotion as STEM fields, which are male-dominated, continue to have higher promotion rates.

**Equal attrition** uses the same gender-shuffle as the equal promotion scenario, but for attrition. This does not make attrition fully gender independent, as attrition rates still increase with years at rank, i.e., time without promotion. As long as promotion is not equitable, attrition will follow gendered patterns.

**Equal hiring rate** bootstraps the hiring-pool with equal women and men. In other words, if our hiring pool originally contains 100 men and 50 women, we bootstrap 50 more women, such that a new hire has a 50% chance of being male/female. Alone, this does not give full equity in hiring. Women are still less likely to be hired into STEM fields and are still more likely to be hired at lower ranks compared to men. The gender differences in age distribution are maintained.

**Total hiring equity** implements the equal hiring rate scenario, but also shuffles all other demographics at random; i.e., there are equal numbers of women and men in the hiring pool and gender is not correlated with any other demographics (but other demographics retain covariance). In this case, men and women are equally likely to be hired into STEM or non-STEM disciplines and at the same ranks at the same age.

Similar to James & Brower [4], we find hiring to be the most effective institutional gender equity intervention (Figure 4). Without intervention, we predict that future women will continue to be underrepresented at all ranks. Lecturers will be 52.4% women; Senior Lecturers 35.1%, Associate Professors 31.4%, and Professors 30.5% (Figure 4). The equal hiring rate intervention (equal women/men in the hiring pool but hiring rank is gender dependent) increases the proportion of women across all ranks by about 10 percentage points (Figure 4). Specifically, women’s representation at the Lecturer rank increases, on average, by 11.2 percentage points to 63.6%. Without a commensurable representation increase at other academic ranks, more female Lecturers may *increase* gender rank gaps. However, at the Senior Lecturer rank, this increase is 11.5 percentage points, and the increase is 10.5 percentage points at the Associate Professor rank and 9.9 percentage points at the Professor rank. This general pattern holds across both STEM and non-STEM disciplines.

**Figure 4:**
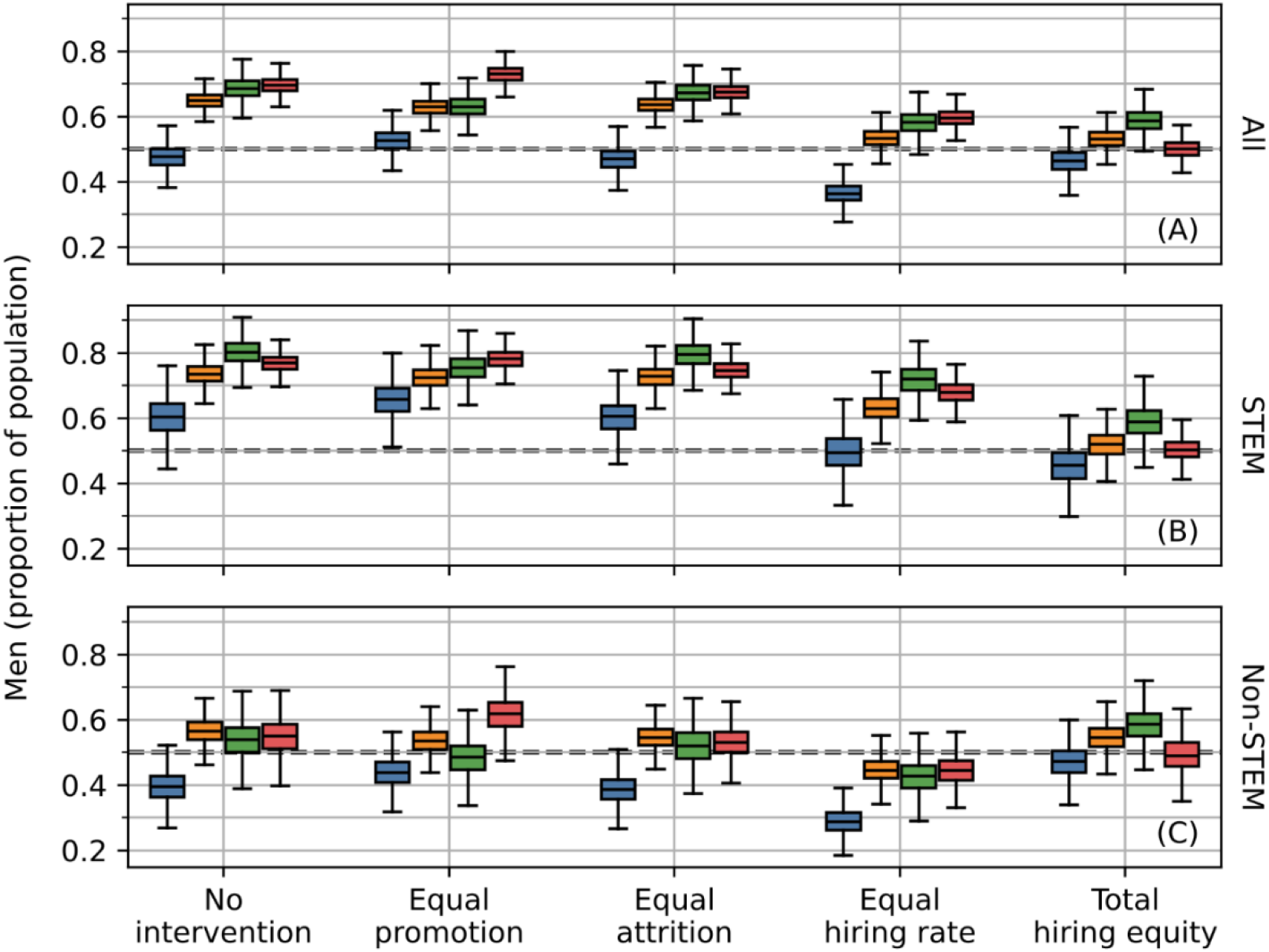
Total hiring equity and equal hiring rates are the best institutional interventions for achieving long-term gender parity across all ranks. Boxplots (median, IQR, 90^th^ percentile) showing the long-term prediction for all institutional interventions, for (A) total, (B) STEM, and (C) non-STEM populations. Equal hiring rates increase the proportion of women at all ranks, whereas total hiring equity achieves near full gender parity at all ranks. Promotion and attrition equality show marginal parity improvements compared to the no-intervention scenario. Blue - Lecturer rank; orange - Senior Lecturer rank; green - Associate Professor rank; red - Professor rank; dotted line - gender parity.

Interventions targeting promotion and attrition are less effective than the equal hiring intervention. The equal promotion intervention increases the number of women Associate Professors by 5.6 percentage points, and the equal attrition intervention increases the number of women Professors by 2.0 percentage points. These are the largest changes in gender composition under each intervention, and they are around half of the smallest change in the equal hiring case.

The total hiring equity intervention (gender equality in the hiring pool and a full demographic shuffle) reduces gender gaps across all ranks (Figure 4). Full gender parity (which we define as the interquartile range (IQR) spanning 50%) is achieved at the Professor rank (50.0% women; IQR: 48.1-51.8%). The Lecturer, Senior Lecturer, and Associate Professor ranks show marked improvement, but, on average, fail to reach full parity with 53.6%, 46.8% and 41.3% women, respectively.

The difference in efficacy between the equal hiring rate and total hiring equity interventions suggests that many gender gaps are widened by other demographic disparities. Therefore, we explore a wide range of intervention strategies by shuffling gender hiring rates, rank at hire, age at hire, and academic discipline. To limit the total parameter space, we assume that academic promotion and attrition are equal with respect to gender. We choose to fix promotion and attrition, as these can be fully determined by top-down institutional management, whereas hiring interventions are partially exogenous.

Simply hiring more women is unlikely to resolve gender representation gaps, as even when women and men are hired at equal rates, with equal promotion and attrition thereafter, men will remain overrepresented at the highest academic rank: Professor (Figure 5). However, hiring women and men at the same rate, with the same rank and discipline distributions (given existing gendered age-structures) closes long-term Professor gaps (ensuring promotion/attrition equity). This claim is supported by the fact that both interventions in our simulation that achieve gender parity target *hiring rate, rank*, and *discipline*. Other interventions, such as equalising age at hire between women and men, are comparatively ineffective, even in conjunction with equal promotion/attrition.

**Figure 5:**
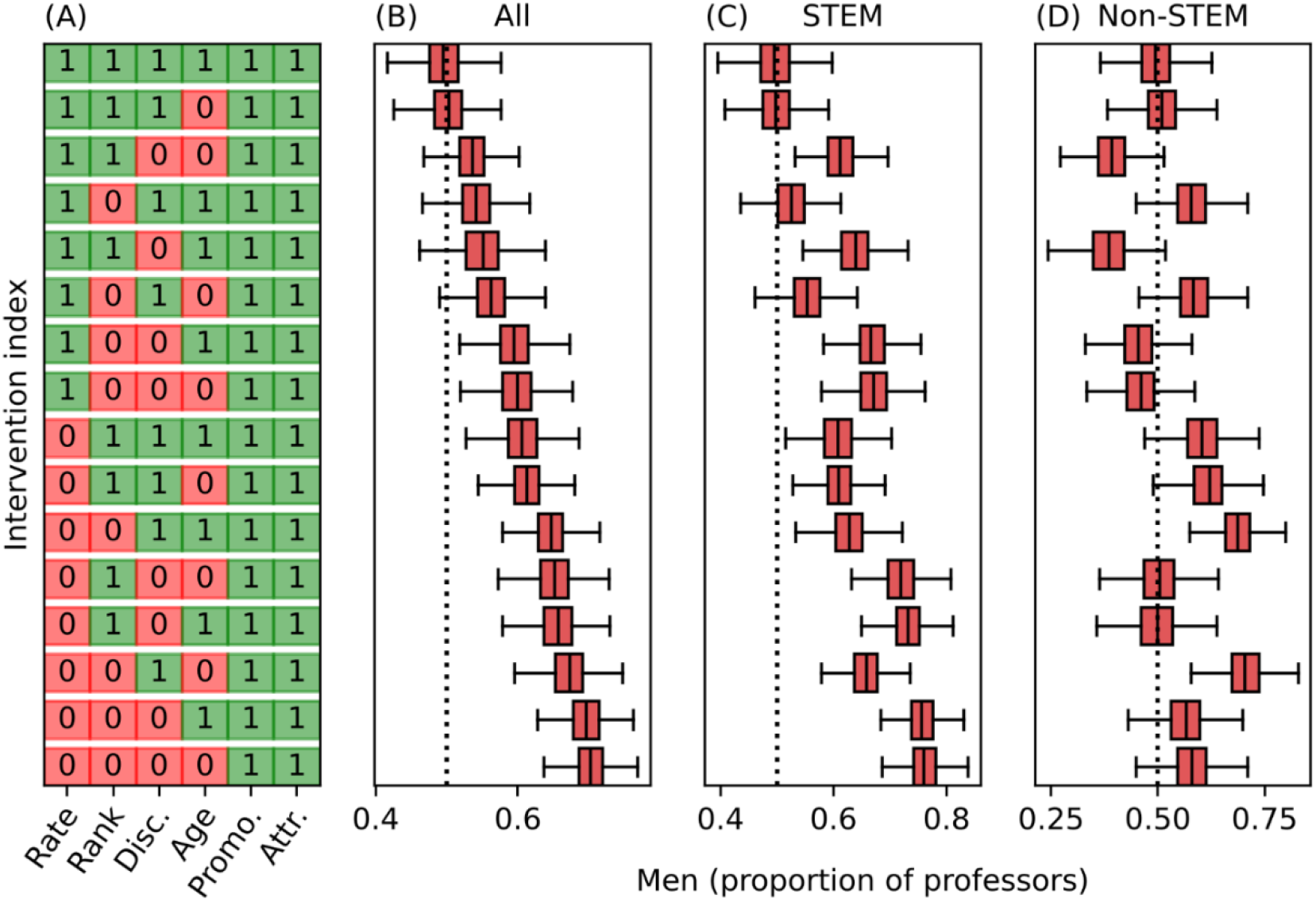
Comparing intervention combinations, full gender parity at the Professor rank is consistently achieved through interventions targeting hiring rate, rank, and discipline. The long- term proportion of male Professors (red boxes; median, IQR, 90^th^ percentile) following institutional interventions for (A) all individuals, (B) STEM faculty and (C) non-STEM faculty. Intervention index denotes the targeted demographics/career progression mechanisms (D) which are coded as: *rate* - equal hiring rate; *rank* - equal rank at hire; *disc*. - academic discipline; *age* - biological age at hire; *promo*. - equal promotion; *attr*. - equal attrition. Interventions maintain covariance between all demographics, other than gender.

Overall gender parity can *almost* be achieved without ensuring equitable academic discipline distributions between women and men. However, disaggregating our population into STEM and non-STEM fields, we find that such scenarios are achieved only via disciplinary segregation, where a lack of female STEM Professors is offset by a preponderance of female non-STEM Professors. Due to the overall higher promotion rates (and thereby ranks) in STEM, women remain underrepresented at the Professor rank en masse.

## 6. Gender pay gaps

By connecting each year within a rank with the associated salary step (see Supplementary Materials for full rank-pay details), we can calculate the average salary across all individuals in the model at any time. We can also track the lifetime pay of any individual from when they were hired to when they leave academia. We define two measures of salary gap: instantaneous and lifetime. The instantaneous gap is the pay gap measured at time *t* using the current pay of all individuals, shown as the difference between the average men’s salary that year, 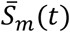, and the average women’s salary, 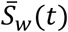, relative to the average men’s salary,

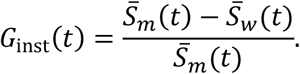

The lifetime gap is the career pay gap between women and men using the whole lifetime pay of each individual. For this measure, we only include individuals who have left the system to ensure they have a full career of data. To make the measure less noisy, we include all individuals who have ended their career in the last 50 years (about 1 academic generation). In other words, this is the difference between the average men’s career pay, 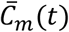, and the average women’s career pay, 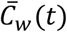, relative to the average men’s career pay, of individuals whose career ended between *t*and *t* −50,

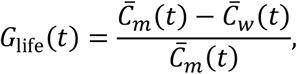

where 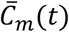is the average lifetime pay of a male academic and 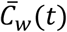is the equivalent for women.

Adjusting our population size, we can simulate an arbitrary department/school in a university. We initialise our simulated population with *N* = 100academic individuals (compared to *N* = 1,000for the full university). Our individual-based model runs for *t*_max_ = 2,500, after which the first 500 and final 100 years are discarded. Academic departments are often drawn around disciplinary boundaries, so we simulate both STEM and non-STEM departments. We assume that women and men are hired at equal rates, but other aspects of hiring, for example, age and rank at hire, follow the distributions from the data set.

Figure 6 shows time-averaged gender representation and pay gaps. In our STEM and non-STEM fields, the median instantaneous gender pay gaps are 5.1% (NZ$7,016) and 4.7% (NZ$6,987), respectively. These are close to the NZ national workforce average of 5.2% [33].

**Figure 6:**
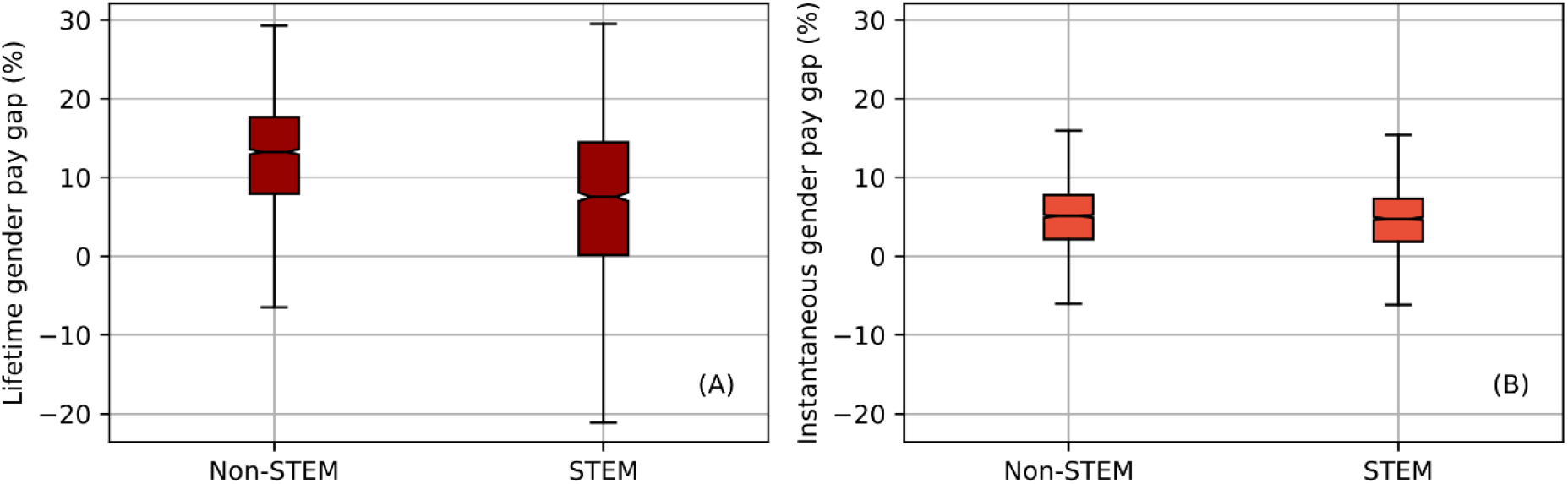
Median lifetime gender pay gaps are 13.3% and 7.6% in non-STEM and STEM departments, respectively, whereas the instantaneous gender pay gaps are both about 5%. The lifetime (A) and instantaneous (B) gender pay gaps (median, IQR, 90th percentile) in STEM and non-STEM departments. Orange - instantaneous gender pay gap; maroon - lifetime gender pay gap.

The instantaneous gender pay gap is often chosen as a measure of inequality [34]. However, we find that measuring gender pay inequity using only instantaneous metrics may underestimate gender pay inequity on the career scale. For example, the median lifetime gender pay gap for the non-STEM department is 13.3% (NZ$270,204). The median lifetime gender pay gap in STEM is smaller at 7.6% (NZ$195,501). The lifetime gender pay gaps have larger variance than the instantaneous gender pay gaps, as they can vary wildly depending on who is hired and who leaves in a given academic cohort. This indicates that even if individual departments were to equalise hiring rates between women and men, there remain large economic disparities. For comparison, Brower & James [2] have estimated the average NZ university has a lifetime gender pay gap of about NZ$400,000.

Both the non-STEM and STEM lifetime gender pay gaps are caused by the same phenomena: more women end their careers at the Lecturer rank than men. However, this early-career disparity affects more academic staff in non-STEM fields, compared to their STEM counterparts. Due to the higher promotion rates in STEM disciplines, academic staff working in STEM spend less time at lower ranks where the largest gender disparities occur. Of course, this is all contingent on an equal gender balance in STEM hiring. What our model indicates is that STEM women who do make it past the Lecturer rank generally tend to be well-promoted. Whether this is the result of *survivorship bias* [4, 35, 36], we cannot say.

### 7. Study limitations

Our data has several limitations. The College of Education institutional restructure in 2007 greatly impacted our data set. It made comparisons between pre-2007 and post-2007 periods nearly impossible, and given that the College of Education is predominantly women, it impaired our ability to infer women’s hiring rates, as over 100 women appear in our data without hire. We approximated annual salary, as the UC payroll data did not contain this information. Although the vast majority of academics are on the TEU Collective Employment Agreement, this ignores several (potentially lucrative) income streams.

A single institution case study has advantages associated with the level of detail available. However, a larger sample size would have allowed for stronger results, especially with rare events, such as Professor hiring. This being said, most UC demographics and hiring practices are similar to other international institutes. For example, data from the USA show that men receive their PhD earlier than women [37] which would likely result in earlier hiring, and most progress in USA academic gender representation was due to older men leaving, rather than present hiring rates practices [38]. Elite women are disadvantaged in USA hiring networks [39], which resembles the lack of women Professors hired at UC. Finally, women tend to be underrepresented in STEM fields internationally [40-42], and many STEM departments receive pay and promotion benefits [43-45].

We do not have data on job applications, i.e., the hiring pool for each job listing. Therefore, we were limited in building statistical models of career progression, only including promotion and attrition; especially when considering that hiring seems to be driving representation gaps. For simplicity, our individual-based model assumed a fixed academic population, but we know our academic population is growing over time. Finally, we used case resampling bootstrapping for adding academics to our simulated population, whether that be to initialise our model or to simulate academic hiring. In the future, when more hiring data becomes available, demographic distributions may be estimated parametrically.

## 8. Discussion

We used models parameterised with 15 years of data from one university to examine gender gaps in pay and representation. We find that career progression is neither uniform nor linear; and this non-linearity takes several forms, creating 5 compounding costs of being female in academia.

### Sticky floors – women get stuck at the lowest rank

The non-linearity means years at rank is important. Past a certain point, the longer a person has stayed at their current rank, the less likely they are to be promoted to a higher rank. For women, this hits the hardest at the Lecturer level, meaning many women never make it beyond Lecturer. This is an effect that has been seen elsewhere, for example, Moss-Racusin et al. [31] found that when young men were employed they were more likely to be given mentoring than the equivalent woman. Studies also show that women are less likely than men to receive sponsorship in their careers [46-48] which could explain men having an early career boost. Another explanation could simply be that the majority of female lecturers are hired between the ages of 30 and 40, which is coincidentally around the same time many women start a family [49-53]. This would add support to the need for good parental leave and return to work packages.

Conversely, at the highest promotion level from Associate Professor to Professor, these gender differences have vanished and in some cases women at this level have an advantage over men. This positive effect at higher levels could be a simple case of survivor bias – women who make it this far are very likely to be high achievers having made it through the “hostile obstacle course” that is an academic career for women [54].

### STEM disciplines, coincidentally male-dominated, promote faster

The negative effects that we see for women’s early promotion are independently compounded by discipline. Being in a STEM field has a significant positive effect on promotion at every level compared to a non-STEM field. STEM fields are also male-dominated (at UC and globally [40, 41]). At UC a quarter (25.3%) of female academics are in STEM fields, while over half (55%) of men employed at UC are in STEM fields. This gives the average man a promotion and pay advantage over the average woman. This effect is very unlikely to be unique to UC. James et al. [55] showed that in over 30 countries, individuals working in male-dominated fields (often but not always STEM-related) were more likely to be successful in grant applications and to receive higher research productivity scores. In short, across 30 countries research in male-dominated fields is judged to be better than research in female-dominated fields. This resembles previous findings that disciplines where it is believed “raw, innate talent” is required for success tend to be male-dominated because it is commonly believed women do not have the requisite talent [56]. Perceptions of research achievement and funding success rates are cornerstones of most academic progression which could result in this STEM effect being widespread.

### Complicated relationship between age, promotion, and gender

Another compounding factor for women is age. We find that the relationship between age, gender, and promotion is complex, but ultimately disadvantages women. As academics age, their chances of promotion decrease. Decreasing promotion chances with age is not a phenomenon unique to either UC, Aotearoa New Zealand, or even universities, as the effect is well-known outside academia [57, 58]. Since women tend to be hired older than men, this is an immediate promotion disadvantage from the outset, irrespective of academic rank and discipline. The early-career promotion boost we find for men, but not women, only adds to this relative disadvantage.

### Women hired at lower rank (and salary) than men

By calculating the probability of ultimate promotion from the Lecturer rank to the Senior Lecturer rank, i.e. the probability of reaching Senior Lecturer before leaving academia, we demonstrated the disadvantage that newly hired female Lecturers experience in promotion. This analysis hides the fact the women are more likely to be hired at Lecturer level than men: 70% of women are hired at the Lecturer rank, compared to only 50% of men. This is despite women being slightly older at hiring which would suggest more previous work experience.

Overall, women are more likely to be hired at a lower rank where they are more likely to get stuck. Women also have the additional hurdle of another promotion application before reaching the professoriate. Gender gaps in first jobs after education, although smaller than later-career gender gaps, are well-documented; for example a German study [59] showed women entered the post-university job market on lower salaries than their male counterparts.

### Attrition and years at rank affect women more

We find the probability of leaving the university, i.e. attrition, is less dependent on gender and discipline than hiring and promotion but it does change with academic rank, although equally for men and women and similarly in STEM and non-STEM fields. However, a key driver of attrition is lack of progression. The more years at rank, the higher the likelihood of attrition before promotion. Although this effect is independent of gender, this hides our finding that women are less likely to be promoted and hence will spend more time at a particular rank; thus increasing their chance of leaving in comparison to men [60, 61]. Our finding that women spend more years at lower ranks means this affects women more.

## 9. Conclusion

We find intriguing differences in hiring, promotion, and attrition patterns between STEM and non-STEM. These disciplinary differences further strengthened our conclusion that gender effects are numerous and compounding. This means hiring interventions alone will not close the gender gaps. We find the most effective combination of interventions equalises gender balance in hiring, closes the gender rank gaps at hire, and narrows the gender representation gap in STEM.

In summary, while limited by their assumptions and data availability, our novel individual-based model harmonises existing literature into a single recommendation. Universities seeking effective gender parity initiatives should hire more women at equal ranks and in similar disciplines to men; just hiring more women will not close the gaps.

## Supporting information

Employment data, inferred salary data, and model selection details

## References

1. Barrett-Walker, T., et al., Stochastic modelling of intersectional pay gaps in universities. Royal Society Open Science, 2023.

2. Brower, A. and A. James, Research performance and age explain less than half of the gender pay gap in New Zealand universities. PLoS ONE, 2020. 15(1): p. 1––13.

3. Hargens, L.L. and J.S. Long, Demographic inertia and women’s representation among faculty in higher education. Journal of Higher Education, 2002.

4. James, A. and A. Brower, Levers of change: using mathematical models to compare gender equity interventions in universities. Royal Society Open Science, 2022. 9(9).

5. Marschke, R., et al., Demographic Inertia Revisited: An Immodest Proposal to Achieve Equitable Gender Representation among Faculty in Higher Education. The Journal of Higher Education, 2007.

6. Shaw, A.K. and D.E. Stanton, Leaks in the pipeline: separating demographic inertia from ongoing gender differences in academia. Proc Biol Sci, 2012. 279(1743): p. 3736–41.

7. Thomas, N.R., D.J. Poole, and J.M. Herbers, Gender in Science and Engineering Faculties: Demographic Inertia Revisited. PLoS One, 2015. 10(10): p. e0139767.

8. Canterbury, U.o. General Staff Collective Employment Agreement. 2024 [cited 2026 04/03/2026].

9. Ovseiko, P.V., et al., Advancing gender equality through the Athena SWAN Charter for Women in Science: an exploratory study of women’s and men’s perceptions. Health Res Policy Syst, 2017. 15(1): p. 12.

10. Canterbury, U.o. UC’s commitment to the SDGs. 2023 [cited 2026 04/03/2026].

11. Mayer, S.J. and J.M.K. Rathmann, How does research productivity relate to gender? Analyzing gender differences for multiple publication dimensions. Scientometrics, 2018. 117(3): p. 1663–1693.

12. Nygaard, L.P., D.W. Aksnes, and F.N. Piro, Identifying gender disparities in research performance: the importance of comparing apples with apples. Higher Education, 2022. 84(5): p. 1127–1142.

13. van den Besselaar, P. and U. Sandstrom, Vicious circles of gender bias, lower positions, and lower performance: Gender differences in scholarly productivity and impact. PLoS One, 2017. 12(8): p. e0183301.

14. Manfredi, S.L.a.S., Balancing Gender in Higher Education. The European Journal ofWomen’s Studies, 2000. 7(33).

15. Benjamin, E., Disparities in the Salaries and Appointments of Academic Women and Men. Academe, 1999. 85(1): p. 60–62.

16. Monroe, K. and W. Chiu, Gender Equality in the Academy: The Pipeline Problem. PS: Political Science & Politics, 2010. 43: p. 303–308.

17. Baker, M., Career confidence and gendered expectations of academic promotion. Journal of Sociology, 2010. 46(3): p. 317–334.

18. Abramo, G., C.A. D’Angelo, and A. Caprasecca, Gender differences in research productivity: A bibliometric analysis of the Italian academic system. Scientometrics, 2009. 79(3): p. 517–539.

19. Costas, R., T.N. van Leeuwen, and M. Bordons, A bibliometric classificatory approach for the study and assessment of research performance at the individual level: The effects of age on productivity and impact. Journal of the American Society for Information Science and Technology, 2010. 61(8): p. 1564–1581.

20. Huang, J., et al., Historical comparison of gender inequality in scientific careers across countries and disciplines. Proc Natl Acad Sci U S A, 2020. 117(9): p. 4609–4616.

21. Sabharwal, M., Comparing Research Productivity Across Disciplines and Career Stages. Journal of Comparative Policy Analysis: Research and Practice, 2013. 15(2): p. 141–163.

22. Gingras, Y., et al., The effects of aging on researchers’ publication and citation patterns. PLoS One, 2008. 3(12): p. e4048.

23. Falagas, M.E., V. Ierodiakonou, and V.G. Alexiou, At what age do biomedical scientists do their best work? FASEB J, 2008. 22(12): p. 4067–70.

24. Rørstad, K. and D.W. Aksnes, Publication rate expressed by age, gender and academic position-A large-scale analysis of Norwegian academic staff. Journal of Informetrics, 2015.

25. Franks, A.M., et al., Rank and Tenure Amongst Faculty at Academic Medical Centers: A Study of More Than 50 Years of Gender Disparities. Acad Med, 2022. 97(7): p. 1038–1048.

26. Shaw, A.K., et al., Differential retention contributes to racial/ ethnic disparity in U.S. academia. PLoS ONE, 2021.

27. LaBerge, N., et al., Gendered hiring and attrition on the path to parity for academic faculty. Elife, 2024. 13.

28. Spoon, K., et al., Gender and retention patterns among U.S. faculty. Sci Adv, 2023. 9(42): p. eadi2205.

29. Cech, E.A. and T.J. Waidzunas, Systemic inequalities for LGBTQ professionals in STEM. Sci Adv, 2021. 7(3).

30. Weissman, J.L., et al., Queer-and trans-inclusive faculty hiring-A call for change. PLoS Biol, 2024. 22(11): p. e3002919.

31. Moss-Racusin, C.A., et al., Science faculty’s subtle gender biases favor male students. Proc Natl Acad Sci U S A, 2012. 109(41): p. 16474–9.

32. Begeny, C.T., et al., In some professions, women have become well represented, yet gender bias persists-Perpetuated by those who think it is not happening. Sci Adv, 2020. 6(26): p. eaba7814.

33. NZ, S. Gender pay gap narrows to lowest on record. 2025 [cited 2026 04/03/26].

34. Coron, C., Measuring the gender pay gap: the complexity of HR metrics. Employee Relations: The International Journal, 2021. 43(5): p. 1194–1213.

35. Llorens, A., et al., Gender bias in academia: A lifetime problem that needs solutions. Neuron, 2021. 109(13): p. 2047–2074.

36. Fontanarrosa, G., et al., Over twenty years of publications in Ecology: Over-contribution of women reveals a new dimension of gender bias. PLoS One, 2024. 19(9): p. e0307813.

37. Report, W.i.A. The Gender Gap in the Time It Takes to Earn a Doctoral Degree. 2024 30 March 2026]; Available from: https://wiareport.com/2024/02/the-gender-gap-in-the-time-it-takes-to-earned-a-doctoral-degree/.

38. Wapman, K.H., et al., Quantifying hierarchy and dynamics in US faculty hiring and retention. Nature, 2022. 610(7930): p. 120–127.

39. Clauset, A., S. Arbesman, and D.B. Larremore, Systematic inequality and hierarchy in faculty hiring networks. Science Advances, 2015. 1(1): p. e1400005.

40. Chavatzia, T., Cracking the code: girls’ and women’s education in science, technology, engineering and mathematics (STEM). 2017, UNESCO: Paris, France. p. 85 p.

41. Wang, M.T. and J.L. Degol, Gender Gap in Science, Technology, Engineering, and Mathematics (STEM): Current Knowledge, Implications for Practice, Policy, and Future Directions. Educ Psychol Rev, 2017. 29(1): p. 119–140.

42. Baker, S.P. and C. Koedel, Diversity trends among faculty in STEM and non-STEM fields at selective public universities in the U.S. from 2016 to 2023. Humanities and Social Sciences Communications, 2024. 11(1).

43. Byrne, Z.S.C., Kelly A., STEM vs. Non-STEM Cultures in a R1 University. Journal of Organizational Psychology, 2020. 20(1).

44. O’Connor, A.M.G., Sandra Wiley; Bowen, Bonnie Sue, Becoming a Professor: an Analysis of Gender on the Promotion of Faculty from Associate to Full Professor. International Journal of Gender, Science and Technology, 2012. 4(1): p. 78–101.

45. Wilcox, V. and M.A.R. Forhad, Reverses in Gender Salary Gaps Among STEM Faculty: Evidence from Mean and Quantile Decompositions. 2023. 23(4): p. 943–979.

46. Cutter, C.M., et al., Gender Differences in Faculty Perceptions of Mentorship and Sponsorship. JAMA Netw Open, 2024. 7(2): p. e2355663.

47. Levine, R.B., et al., “It’s a Little Different for Men”-Sponsorship and Gender in Academic Medicine: a Qualitative Study. J Gen Intern Med, 2021. 36(1): p. 1–8.

48. Shakil, S. and R.F. Redberg, Gender Disparities in Sponsorship—How They Perpetuate the Glass Ceiling. JAMA Internal Medicine, 2017. 177(4): p. 582–582.

49. Kwon, D., Motherhood derails women’s academic careers — these data reveal how and why, in Nature News. 2026.

50. Tower, L.E. and M. Latimer, Cumulative Disadvantage: Effects of Early Career Childcare Issues on Faculty Research Travel. Affilia-Journal of Women and Social Work, 2016.

51. Antecol, H., K. Bedard, and J. Stearns, Equal but inequitable: Who benefits from gender-neutral tenure clock stopping policies? 2018.

52. Misra, J., J.H. Lundquist, and A. Templer, Gender, Work Time, and Care Responsibilities Among Faculty. Sociological Forum, 2012.

53. Mason, M.A., N.H. Wolfinger, and M. Goulden, Do Babies Matter? 2019.

54. Berhe, A.A., et al., Scientists from historically excluded groups face a hostile obstacle course. Nature Geoscience, 2021. 15(1): p. 2–4.

55. James, A., et al., Female-dominated disciplines have lower evaluated research quality and funding success rates, for men and women. Elife, 2024. 13: p. RP97613.

56. Leslie, S.-J., et al., Expectations of brilliance underlie gender distributions across academic disciplines. Science, 2015. 347(6219): p. 262–265.

57. Machado, C.S.P. Miguel, Age and Opportunities for Promotion. 2013, Institute of Labor Economics (IZA).

58. Hemsley, S. Employers still guilty of overlooking older workers for promotions. 2022 31 March 2026]; Available from: https://www.worklife.news/dei/employers-still-guilty-of-overlooking-older-workers-for-promotions/.

59. Sandner, M. and I. Yükselen, Unraveling the gender wage gap: Exploring early career patterns among university graduates. Scottish Journal of Political Economy, 2025. 72(2): p. e12405.

60. Hunt, J., Why do Women Leave Science and Engineering? ILR Review, 2016. 69(1): p. 199–226.

61. Lowenstein, S.R., G. Fernandez, and L.A. Crane, Medical school faculty discontent: prevalence and predictors of intent to leave academic careers. BMC Medical Education, 2007. 7(1): p. 37.

