## Supplementary material for "The compounding costs of being female in academia: Individual-based modelling of career progression and interventions": Employment data, inferred salary data, and model selection details

### Supplementary Materials


1. Employment data
    - *Hiring rate by gender*
    - *Hiring rank by gender*
    - *Hiring age by gender*
    - *Years at rank by gender*
2. Inferred salary data
3. Model selection
4. References

### 1. Employment data

All staff present in the data are employed on continuing contracts (analogous to tenure in the US). Payroll entries are mostly due to pay increases, which are often annual. Missing years are easily interpolated, assuming fixed demographics (e.g., sex) remain constant, and time-variant demographics (e.g., age) are incremented annually. This yields one record per academic per year. If an academic has more than one data point in a single year, we take only the latest entry. Although Human Resources keeps records on administrative-only faculty, their career progression is typically different from research-intensive faculty, so we exclude them from our analysis.

UC payroll data does not explicitly record new academic hires. However, given the unique institutional ID numbers present in the UC data set, a record containing a new ID number implies a new hire. A hiring anomaly occurs in the year 2007, where, due to institutional restructuring, the College of Education (one of the five colleges at UC) was created. This appears in the data set as $N = 115$ new faculty. We exclude these faculty from our hiring pool. This approach will unintentionally remove some faculty who were hired into the College of Education in 2007, but since we cannot distinguish these individuals from continuing faculty who were annexed into the University, we choose to exclude them all. This will marginally underestimate the number of non-STEM women in the University hiring pool.

Following other NZ universities, the University of Canterbury has four main academic ranks: Lecturer (L), Senior Lecturer (SL), Associate Professor (AP), and Professor (P). Other positions exist, such as Assistant Lecturers and postdoctoral researchers. However, the UC data set only contains records for the four main ranks (L, SL, AP, P). Within each rank, there are a given number of steps. Lecturers and Senior Lecturers typically increase one step per year, whereas Associate Professors and Professors apply for most step-increases (Table S2). Annual pay increases with each step and between each rank. Academic ranks are public via the University website, whereas step is private. For promotion between ranks, an application is required, where academics must demonstrate excellence in research, teaching, and/or administrative work. Double-promotions (e.g., from Lecturer to Associate Professor) and demotions (e.g., from Professor to Associate Professor) are absent from our data. The University of Canterbury Collective Agreement, which dictates academic salary and rank scales, has been reproduced below for convenience (Table S2). We standardise salary comparison by using the 2022 pay scale.

| | | | Employed | | | | Hired | | | |
|---|---|---|---|---|---|---|---|---|---|---|
| **Years** | **Discipline** | **Rank** | *Avg./year* | | *Avg. age* | | *Avg./year* | | *Avg. age* | |
| | | | **W** | **M** | **W** | **M** | **W** | **M** | **W** | **M** |
| 2005-2009 | STEM | L | 11.0 | 19.0 | 38.6 | 35.6 | 3.6 | 6.2 | 38.1 | 33.2 |
| | | SL | 20.8 | 87.6 | 43.2 | 47.0 | 1.2 | 5.2 | 36.3 | 40.3 |
| | | AP | 6.6 | 54.4 | 47.9 | 52.5 | 0.0 | 1.6 | - | 49.8 |
| | | P | 2.0 | 33.8 | 48.3 | 57.7 | 0.0 | 2.0 | - | 56.5 |
| | Non-STEM | L | 68.8 | 43.8 | 45.3 | 44.1 | 8.4 | 6.4 | 39.2 | 39.1 |
| | | SL | 69.6 | 85.4 | 50.6 | 50.9 | 1.0 | 4.8 | 45.2 | 47.7 |
| | | AP | 13.6 | 23.8 | 54.5 | 53.6 | 0.8 | 0.0 | 52.8 | - |
| | | P | 4.0 | 23.6 | 54.6 | 57.4 | 0.6 | 2.8 | 54.3 | 51.3 |
| 2010-2014 | STEM | L | 12.2 | 25.4 | 38.2 | 36.5 | 2.4 | 6.8 | 34.9 | 36.0 |
| | | SL | 25.0 | 80.4 | 43.0 | 46.0 | 0.4 | 2.2 | - | 46.3 |
| | | AP | 9.0 | 38.8 | 47.9 | 51.7 | 0.0 | 0.0 | - | - |
| | | P | 7.6 | 49.8 | 51.0 | 57.5 | 0.2 | 0.0 | - | - |
| | Non-STEM | L | 57.2 | 40.2 | 47.9 | 46.3 | 3.6 | 4.0 | 40.1 | 37.6 |
| | | SL | 70.8 | 72.0 | 51.4 | 51.6 | 1.6 | 1.2 | 46.9 | 42.5 |
| | | AP | 25.0 | 33.8 | 55.1 | 53.5 | 0.4 | 0.4 | - | - |
| | | P | 10.6 | 26.6 | 55.2 | 57.8 | 0.6 | 0.6 | 51.7 | 54.0 |
| 2015-2020 | STEM | L | 15.3 | 23.8 | 37.7 | 38.5 | 5.0 | 8.0 | 36.6 | 37.0 |
| | | SL | 21.7 | 52.7 | 45.0 | 44.9 | 0.8 | 3.5 | 50.8 | 42.2 |
| | | AP | 13.2 | 54.5 | 48.0 | 50.0 | 0.2 | 1.3 | - | 44.2 |
| | | P | 13.8 | 64.2 | 54.0 | 55.9 | 0.0 | 2.0 | - | 47.2 |
| | Non-STEM | L | 37.0 | 28.3 | 47.2 | 47.8 | 5.8 | 3.3 | 42.2 | 38.5 |
| | | SL | 48.7 | 52.5 | 51.2 | 50.3 | 2.2 | 2.0 | 44.1 | 43.2 |
| | | AP | 29.8 | 31.2 | 54.7 | 52.1 | 2.0 | 1.0 | 49.2 | 47.5 |
| | | P | 17.8 | 25.8 | 56.6 | 58.5 | 0.2 | 0.7 | - | 53.0 |

Table S1: **Representation and demographics of women and men hired and employed by the University of Canterbury**. Individual records are aggregated into 5-year bins (except data from 2015-2020, which has a 6-year bin). Cells with less than 0.6 academic staff in the range (3 individuals over 5 years) have redacted age data due to privacy concerns; this includes cells without any data (e.g., no STEM Associate Professors were hired between 2010 and 2015).

During the data period there were 5 main research colleges at the University of Canterbury: the College of Science, the College of Engineering, the College of Business and Law, the College of Arts, and the College of Education. The College of Education was created in an institutional restructuring, which occurred in 2007. Due to data limitations, we choose to binarise academic discipline into STEM and non-STEM. STEM faculty belong to the College of Science or the College of Engineering, whereas non-STEM faculty belong Business/Law, Arts, or Education. Although reductive, there is a large literature on gender dynamics in STEM fields [1-4].

We provide a comprehensive summary of the UC data set in Table S1 above. Together with Figure S5, these provide sufficient data for reproducibility, without violating data privacy. Table S1 can be terse to parse, so we provide several figures below which summarise Table S1 further, highlighting important data features.

*Hiring rate by gender*

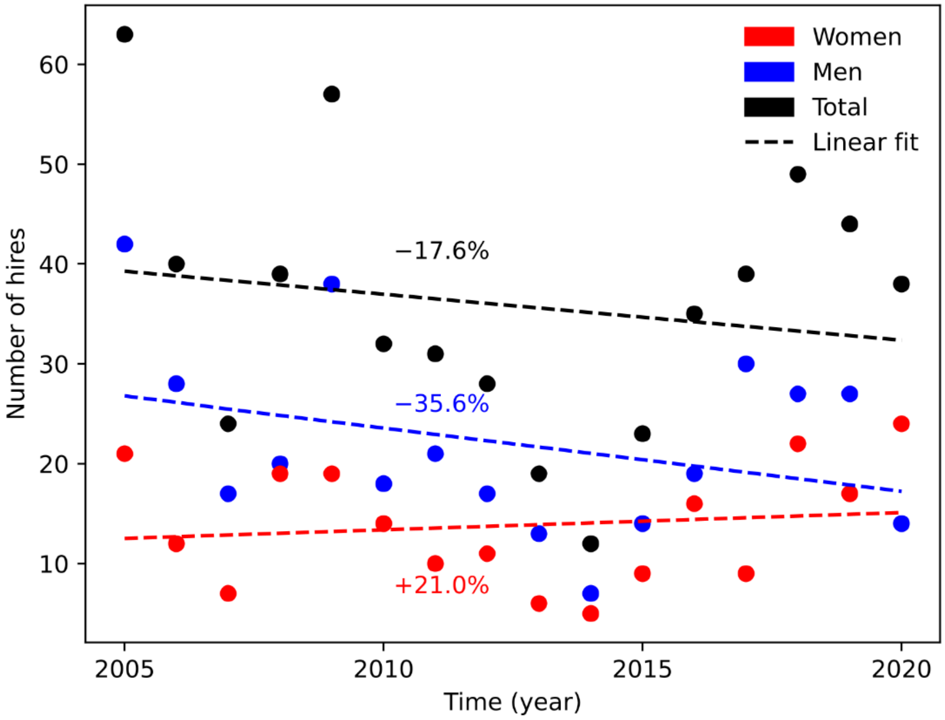


Figure S1: **Over our 15-year data set, women's hiring has increased while men's has decreased.** Percentage change calculated using slope of linear best fit line. Dashed lines – linear best fit; dots – UC data set.

The average number of women hired per year has increased (+21.0%) over the last 15 years (Figure S1). However, this is offset by a decrease in men's hiring (-35.6%), resulting in a net decrease of 17.6%. The primary driver of this decrease was the 2011 Christchurch Earthquake which led to a reduced number of new hires over the 3 years subsequent. To a lesser extent, the College of Education annexation and the exclusion of new College of Education hires deflated the hiring rate in 2007.

*Hiring rank by gender*

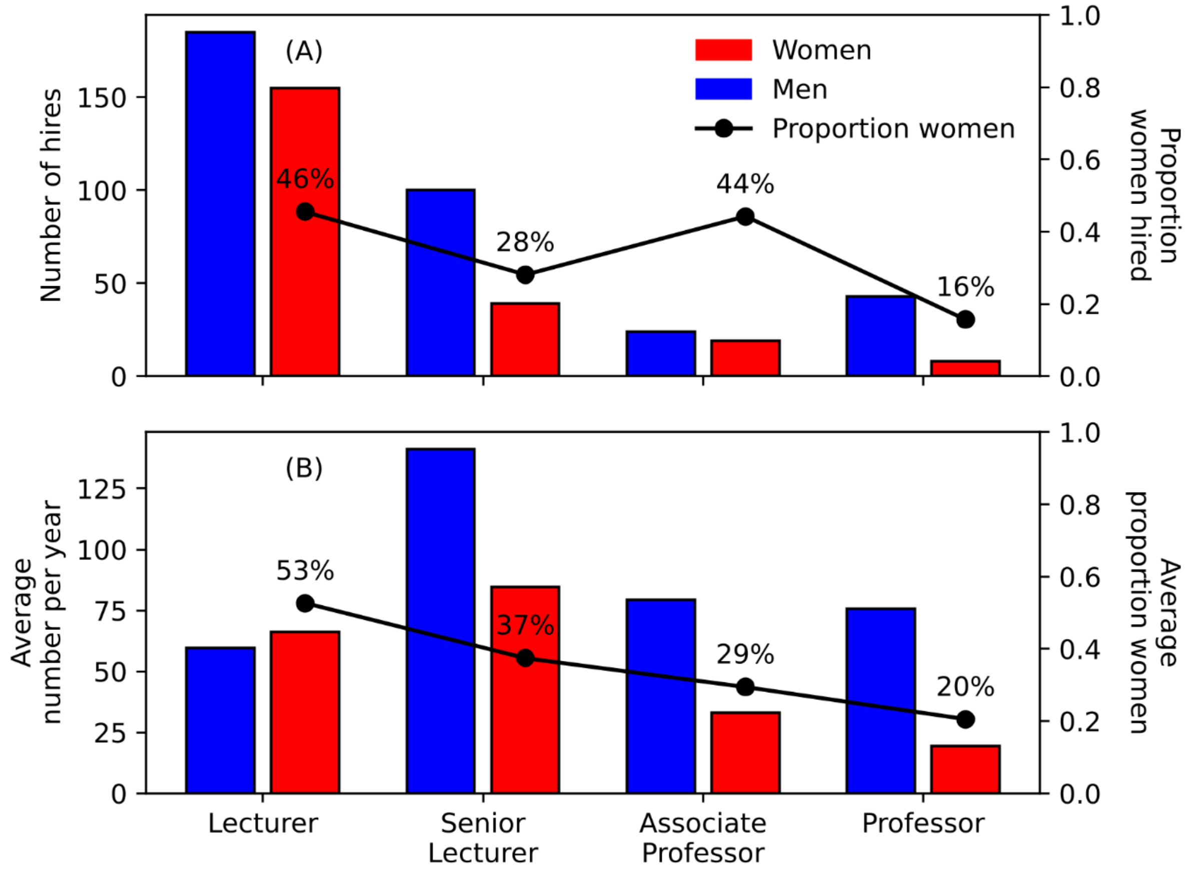


Figure S2: **Fewer women tend to be employed and hired at higher academic ranks compared to men.** The average number of women and men hired at each academic rank (A) and the average number of women and men employed at a given academic rank (B) between 2005-2020 in the UC data set.

On average, the Lecturer rank is comprised of 52.6% women and 47.4% men (Figure S2). In other words, there is a good gender balance of Lecturers. However, with increasing academic rank, the gender balance tends to favour men. At the Senior Lecturer rank, only 37.5% of academic staff are women; at Associate Professor this decreases to 29.4%; only 20.5% of Professors are women. This pattern is generally reproduced in the hiring data. Newly hired Lecturers are 45.6% women. This decreases to 28.1% for newly hired Senior Lecturers; increases again to 44.2% for Associate Professors; decreases to only 15.7% at the Professor rank.

Newly hired Lecturers tend to be about 50% women and 50% men. However, the proportion of women hired into the Lecturer rank tends to be higher than the proportion of men. On average, 70.1% of women - are hired into the Lecturer rank, whereas this number is 52.6% for men. In general, women tend to be hired at lower ranks compared to men with 17.6% of women hired at Senior Lecturer, 8.6% of women hired at Associate Professor, and only 3.6% of women hired at the Professor rank. For men, these numbers are 28.4% at Senior Lecturer, 6.8% at Associate Professor, and 12.2% at Professor.

*Hiring age by gender*

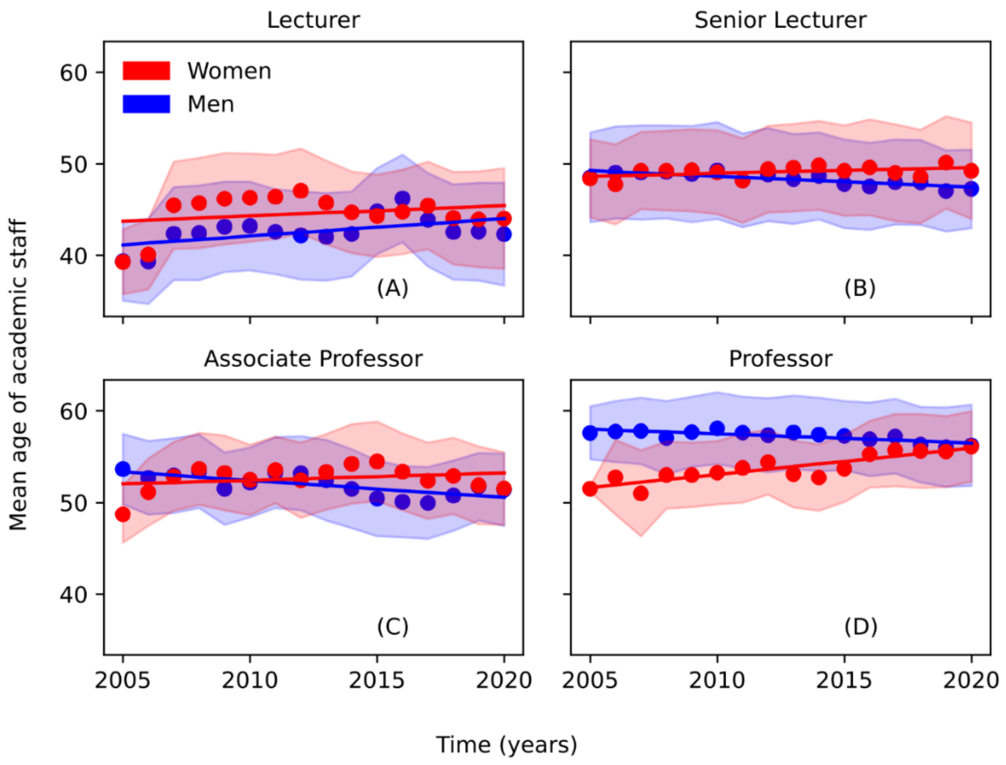


Figure S3: **The average women's age has increased between 2005-2020, while the average man's average has decreased over the same time period.** Mean age of academic staff disaggregated by gender (red – women; blue – men) and rank (A-D). Dots – UC data set; lines – linear best fit; shaded region – standard deviation.

Over our data set (2005-2020), the age profiles of women and men at each academic rank have changed (Figure S3). At the Lecturer rank, both women and men have increased in age. In 2005, the average male Lecturer was 39.4 years old, and in 2020, this increased to 42.3 years old. For women, this increase was from 39.3 years old to 44.0 years old. For the remaining 3 ranks (Senior Lecturer, Associate Professor, and Professor), the average woman tends to be older, while the average man tends to be younger. At the Senior Lecturer rank, the average female age was 48.4 years old in 2005 and increased to 49.2 years old in 2020. This pattern is reversed for men, with an average age of 48.5 years old in 2005 and 47.3 years old in 2020. At Associate Professor, the average woman went from 48.7 to 51.5 years old, and the average man from 53.6 to 51.4. The Professor rank had the largest average female age difference from 51.5 years old in 2005 to 56.1 years old in 2020. Following the pattern established in the previous 2 ranks, the average male Professor tends to decrease in age from 57.6 years old to 56.2 years old.

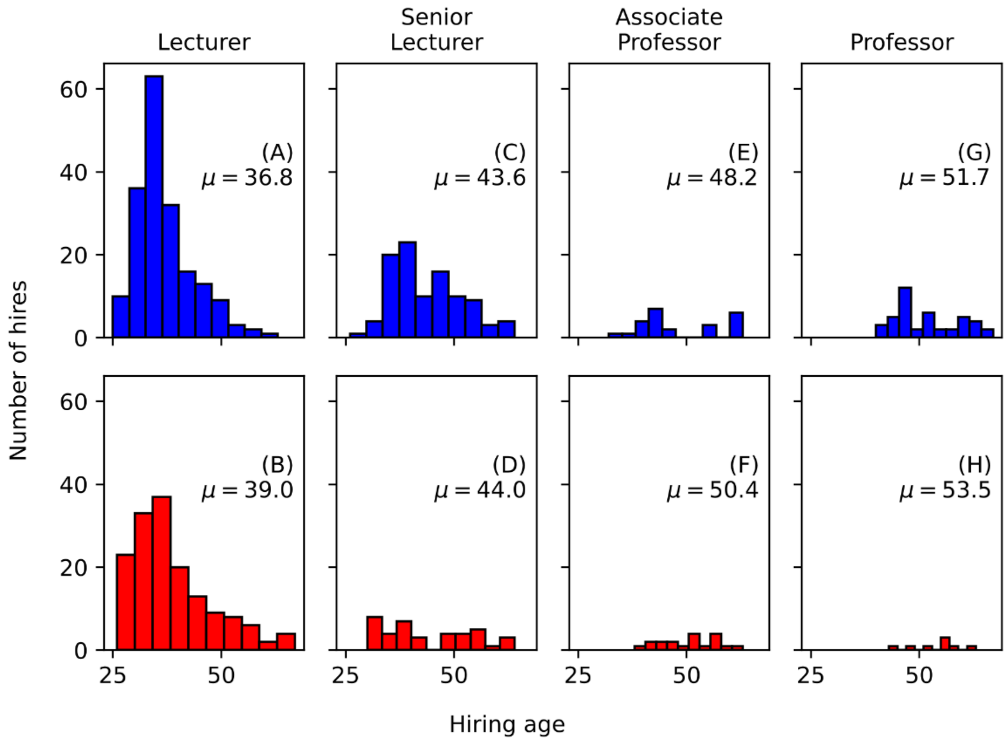


Figure S4: **Women tend to be hired marginally older than men, regardless of academic rank; although this disparity is largest for new Lecturers.** Number of hires between 2005-2020 across all academic ranks and genders (A-H). Red bars – women; blue bars – men.

The average hiring age of women is older than the average hiring age of men (Figure S4). While marginal, this is persistent across all academic ranks. At Lecturer, the average woman hired tends to be 39 years old, whereas the average man tends to be 36.8. Interestingly, women have a larger variance in their hiring age at Lecturer ($\sigma^2 = 77.7$) compared to men ($\sigma^2 = 44.8$). This indicates that not only are women older on average, but they tend to be overrepresented in older academics being hired (e.g., new hires after the age of 50). Since hiring mainly occurs at the Lecturer rank, other ranks do not show such clear patterns. At Senior Lecturer, women tend to be hired at 44 years old; this is 43.6 for men. At Associate Professor, the average woman is hired at 50.4 years old, whereas the average man is hired at 48.2. Although it is rare, the average female Professor woman is hired at age 53.5, and this is 51.7 years old for men.

*Years at rank by gender*

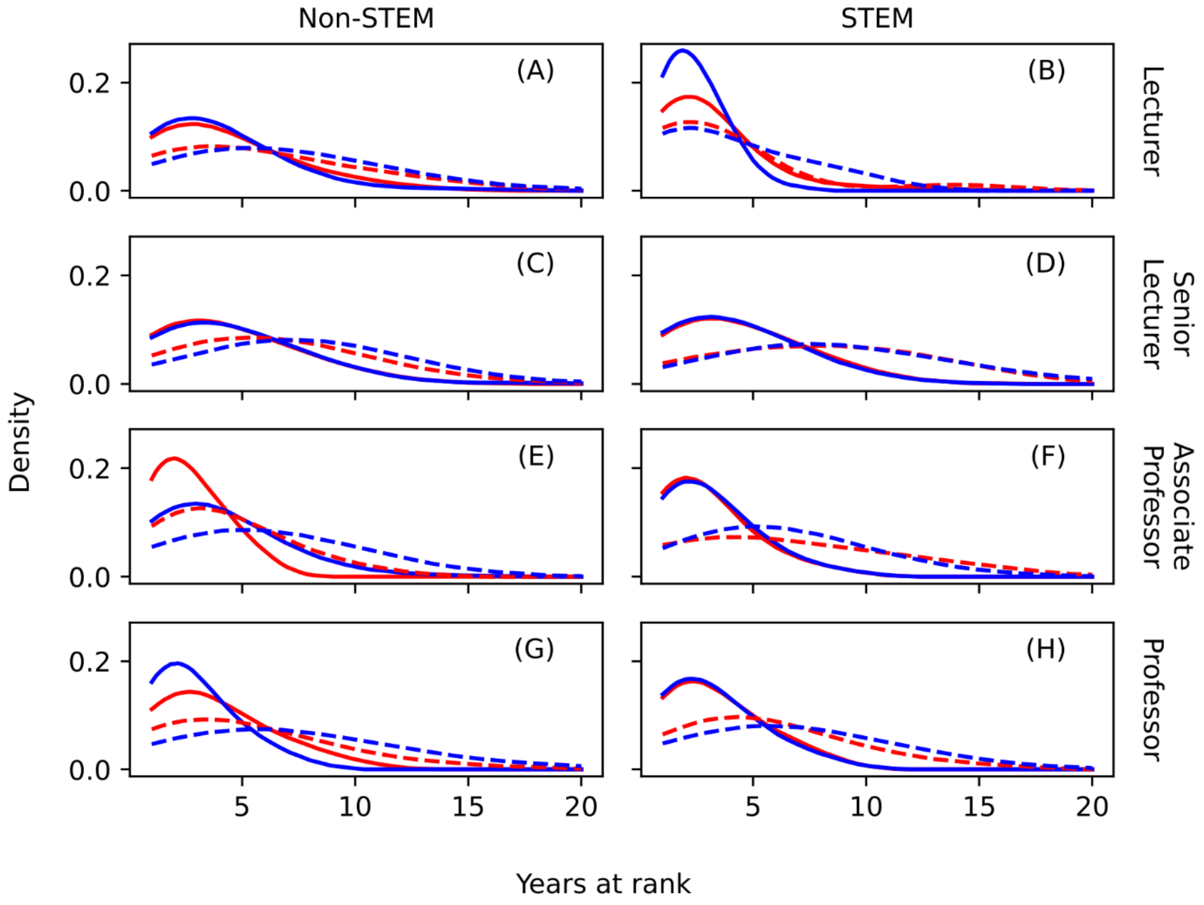


Figure S5: **Men tend to spend longer at higher academic ranks, while women are overrepresented in academics who spend their careers at lower ranks in STEM fields.** Gaussian kernel density estimates of years at rank for STEM and non-STEM disciplines, and all academic ranks. Smoothing kernels here provide anonymity to otherwise identifiable data. Solid lines – academics aged 25-50; dashed lines – academics aged 50-75; red – women; blue – men.

In general, older academic staff (between ages 50-75) tend to spend more years at a given rank than younger academic staff (between ages 25-50) (Figure S5). This is not surprising, since many academics reach their own personal career rank in their 40s or 50s where they spend the rest of their career. Additionally, the variance in years at rank is higher for older staff as older academics have more varied career paths than younger academics.

In STEM fields, younger women have over 3 times the variance ($\sigma^2 = 5.18$) in the number of years spent at the Lecturer rank compared to men ($\sigma^2 = 1.60$), despite having the similar means (2.89 years for women and 2.34 years for men). This suggests that women are overrepresented in career Lecturers, i.e., academics who start and end their career at the Lecturer rank. This pattern is repeated in the 50-75 age bracket, although to a lesser extent: women's mean is 3.23 years with variance $\sigma^2 = 11.1$ and men's mean is 3.79 years with variance $\sigma^2 = 8.27$. Older men tend to spend about one more year at the Professor rank (6.67 year mean; 15.2 year-squared variance), compared to women (5.45 year mean; 11.9 year-squared variance). This extra year spent at Professor at the end of an academics' career will contribute to the lifetime gender pay gap, i.e., the average male Professor will receive one additional salary payment than the average female Professor.

In non-STEM fields, older men tend to spend 2 additional years at the Associate Professor and Professor ranks (6.10 mean years at AP; 6.96 mean years at P) compared to women (4.14 mean years at AP; 4.99 mean years at P). Interestingly, compared to young men, young women spend more time at the Professor rank (women: 3.66 year mean; men: 2.81 year mean), but less time at the Associate Professor rank (women: 2.60 year mean; men: 3.84 year mean).

### 2. Inferred salary data

Academic salaries are dictated by the Tertiary Education Union (TEU) Collective Employment Agreement [5] which maps academic rank and step to annual salary in NZD. For example, in 2022, a 3rd year Lecturer, i.e., a Lecturer on step 3 will make NZ$90,343 before tax; a Professor on step 3 will make NZ$172,587 before tax. We have annual rank and step data for each UC academic, so we can retrieve academic salary. When step data is absent, e.g. when predicting future populations, we approximate salary, $S$, within each rank, $R$, using the number of years spent at a given rank, $\tau$. We use the model

$$S_R(\tau) = \alpha_R + \beta_R\left(1 - e^{-k_R\tau}\right),$$

where $S_R(\tau)$ is the predicted academic salary after $\tau$ years at rank $R$, $\alpha_R$ is the minimum salary within each rank $R$, $\beta_R$ is the difference between the minimum salary and the maximum salary within each rank $R$, and

$k_R$ is the rate of salary increase within each rank $R$. We fit our model to UC data using the Levenberg-Marquardt algorithm to solve the non-linear least-squares optimisation problem. This model fit accurately replicates the mean academic salary at each rank; although achieves middling $R^2$-values, due to the high variance in individual academic careers (Figure S6).

Nominal rank sometimes does not match pay rank. Academics entering new disciplines may retain their nominal title (e.g. Professor) but are paid as an Associate Professor. This is the case for $N = 19$ staff in our data set. When inferring salary, we use pay rank as a predictor. In our individual-based model, we will assume that academics' nominal rank matches their pay rank.

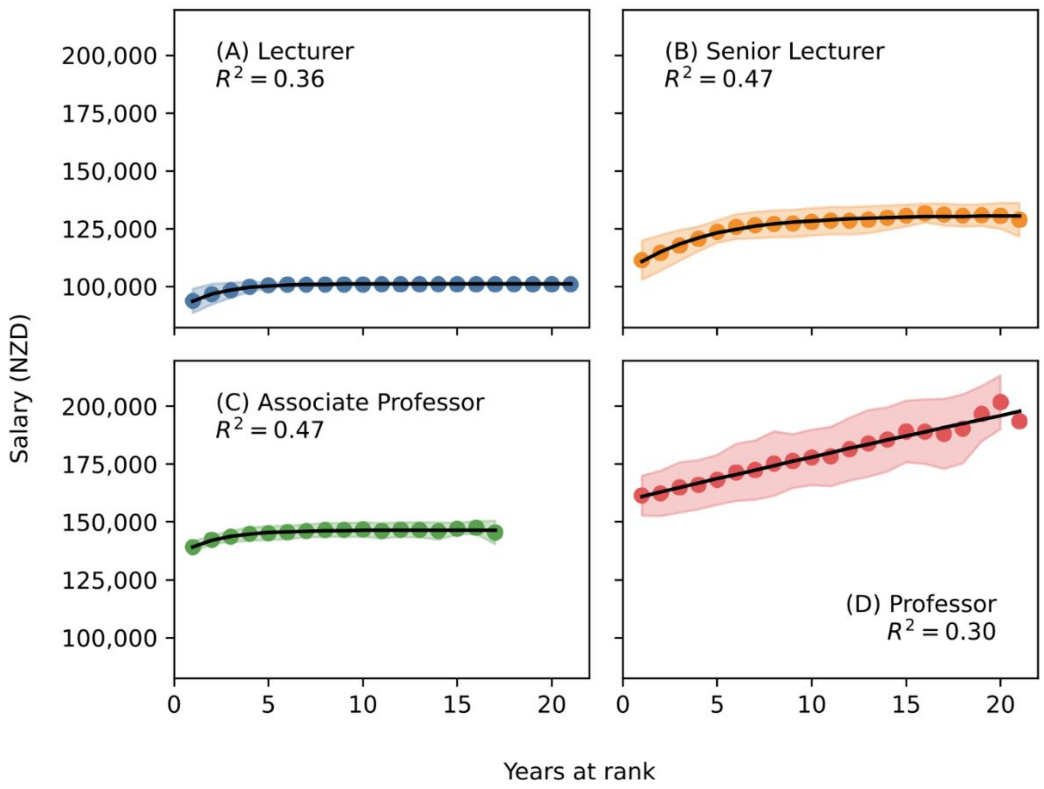


Figure S6: **Our simple model of salary progression fits the average academic, but scores low on Pearson's $R$-squared due to high variance in the data.** The salary model (black line) is fit to UC data (mean academic salary – dots; standard deviation – shaded region) using non-linear least squares. Measuring $R^2$ between our salary model and the data mean achieves very high scores ($R^2 > 0.95$), but scores drop when measured using the full data set indicating high variance around the mean. Blue – Lecturer rank; orange – Senior Lecturer rank; green – Associate Professor rank; red – Professor rank.

| Academic rank | Step | Criteria | Salary (NZD) | Salary (USD) |
|---|---|---|---|---|
| Lecturer (L) | 1 | *Promotion* | $82,882 | $53,459 |
| | 2 | *Automatic* | $86,695 | $55,918 |
| | 3 | *Automatic* | $90,343 | $58,271 |
| | 4 | *Automatic* | $93,825 | $60,517 |
| | 5 | *Automatic* | $97,305 | $62,762 |
| | 6 | *Automatic* | $100,954 | $65,115 |
| Senior Lecturer (SL) | 1 | *Promotion* | $106,258 | $68,536 |
| | 2 | *Automatic* | $109,902 | $70,887 |
| | 3 | *Automatic* | $113,551 | $73,240 |
| | 4 | *Automatic* | $116,867 | $75,379 |
| | 5 | *Automatic* | $120,514 | $77,732 |
| | 6 | *Promotion* | $125,819 | $81,153 |
| | 7 | *Automatic* | $129,467 | $83,506 |
| | 8 | *Automatic* | $133,110 | $85,856 |
| Associate Professor (AP) | 1 | *Promotion* | $138,251 | $89,172 |
| | 2 | *Automatic* | $141,897 | $91,524 |
| | 3 | *Progression* | $145,543 | $93,875 |
| | 4 | *Progression* | $149,026 | $96,122 |
| Professor (P) | 1 | *Promotion* | $158,390 | $102,162 |
| | 2 | *Progression* | $164,962 | $106,400 |
| | 3 | *Progression* | $172,587 | $111,319 |
| | 4 | *Progression* | $179,724 | $115,922 |
| | 5 | *Progression* | $186,651 | $120,390 |
| | 6 | *Progression* | $193,552 | $124,841 |
| | 7 | *Progression* | $199,137 | $128,443 |
| | 8 | *Promotion* | $210,013 | $135,458 |

Table S2: **University of Canterbury Collective Employment Agreement (2022)**. Academic salary and rank information for each continuing research position at UC [5]. Promotions are required for transition between academic ranks (and important within-rank transitions); progressions are required for step-increases at higher ranks; automatic transitions occur annually at lower ranks. NZD to USD conversion rate is the 10-year average between 2017-2026 which is 1 NZD = 0.645 USD. Recent Agreements are available via the University website.

Academic staff often have additional income on top of their University salary. Many academics freelance private consulting work, receive residuals for patents, and own rental property. This is outside of our research scope.

Both the instantaneous and lifetime UC gender pay gaps have decreased since 2007, but not significantly since 2010. There appears to be an increase in gender inequality in 2007, likely due to the College of Education restructure. We assume that the University has always had gender pay gaps around the 2007 value, but that this was hidden by the previous college structure. In other words, gender gaps were not caused, but rather unveiled, by the College of Education annexation. Using linear regression analysis on the gender pay gaps, we find that no significant progress has been made since 2010 ($p > 0.05$; year coefficient on left-censored data).

### 3. Model selection

We use generalised linear models (GLMs) to estimate the correlation and statistical significance between demographics, such as age and gender, and promotion and attrition. Since promotion and attrition are binary events (can be coded using only values 0 and 1), we use logistic regression with a logit link function. Our model fitting process aims to be mathematically rigorous, while avoiding parameter combinations which are not sociologically justified. We begin with three key elements of promotion and attrition, and then select composite models based on statistical goodness of fit and model parsimony. For completeness, each composite model is fit with a time coefficient; this coefficient is insignificant unless stated otherwise.

Previous research has noticed that early-career men may receive more support and mentorship than early-career women [6]. Explanations for this discrepancy vary, but regardless of the cause, the effect is prominent. We hypothesise that, due to less support and mentorship, young women may have lower promotion rates and higher attrition rates than young men. This is operationalised as interaction terms between age and gender and rank and gender.

The University of Canterbury has one of the largest engineering schools in New Zealand, so it is unsurprising that many institutional resources are dedicated to STEM subjects. We measure the impact that binarised discipline (STEM/non-STEM) has on promotion and attrition rates with the models.

Gender gaps in academia are often explained using biological age [7-9]: on average, male researchers tend to be older than female researchers, and older researchers tend to publish more [9-14]. Previous research at the University of Canterbury has already found that biological age explains far less of the academic gender pay gap than expected if age is the main driver [15]. We add nuance to this argument by disaggregating 'age' into three different components: biological age, years at rank, and retirement eligibility. The *years at* *rank* variable is the number of years spent at a given rank before promotion/attrition. We expect years at rank to have a non-linear relationship with promotion/attrition. Most academics spend several years at an academic rank before promotion/attrition, but many years spent at a given rank indicates that an academic may have hit their career rank, i.e., their own personal academic ceiling. Retirement age is 65 years old in New Zealand, and although retirement is not compulsory, we expect academics to leave more and promote less after reaching this age.

We identify which of these three promotion/attrition effects are most prominent at the University of Canterbury by using pairwise model comparison. Models were fit for individual effects, all linear two-way combinations, and a full, three-way model. To ensure that we did not exclude important gender effects, a saturated model was fit, with interactions between gender and all other variables. A final refinement stage included manual backward stepwise regression on each selected model (which marginally increased model parsimony). We selected models using AIC (Table S3), BIC gave the same choices, *k*-fold cross-validation showed no signs of overfitting. We select the promotion and attrition models,

$$\text{logit}(Promotion) \sim IsMale * (Rank + Age) + Rank * YearsAtRank^2 + IsSTEM + Retirement$$

$$\text{logit}(Attrition) \sim IsMale * Age + Rank * YearsAtRank^2 + Retirement.$$

Considering each model fit with a time coefficient, there is some evidence to suggest that STEM promotion rates have increased in recent years (6% annual odds ratio increase; $p = 7.46 \times 10^{-3}$). Given our overall findings, this makes our work timely. Moreover, the average number of years spent a given academic rank may be increasing. However, this is likely due, at least in part, to the left-censoring of our data set. In general, time effects only reach statistical significant in a handful of interactions, and may be alternatively attributable to Type-I error.

Using our selected models, we can calculate the probability of ultimate promotion, $p_U$. This is the probability that a newly hired Lecturer is promoted to Senior Lecturer before the end of their career. To compute this probability, we perform the following calculation,

$$p_U = \sum_{t=0}^{\infty} p_P(t)\big(1 - p_A(t)\big) \prod_{\tau=0}^{t} \big(1 - p_P(\tau)\big)\big(1 - p_A(\tau)\big),$$

where $p_U$ is the probability of ultimate promotion, $p_P(t)$ is the probability of promotion from Lecturer to Senior Lecturer at time $t$, and $p_A(t)$ is the probability of attrition at time $t$. We consider time to be discrete ($t = 0,1,2, \ldots, t_{\max}$) since our ABM runs on discrete time. For simplicity, we are suppressing the dependence of promotion/attrition on age, rank, gender, years at rank, retirement, and discipline. However, given an initial age and years at rank, both are fully described by time, $t$.

| Response (logit) | Predictors (in Wilkinson notation) | $k$ | AIC | BIC |
|---|---|---|---|---|
| ***Promotion*** | ***IsMale * (Age + Rank) + Rank * YAR$^2$ + Ret + IsSTEM*** | **16** | **3904.3** | **4014.9** |
| *Promotion* | *IsMale * (Age + Rank * YAR$^2$ + Ret + IsSTEM)* | 24 | 3906.1 | 4072.0 |
| *Attrition* | *IsMale * (Age + Rank) + Rank * YAR$^2$ + Ret + IsSTEM* | 20 | 3958.5 | 4100.4 |
| ***Attrition*** | ***IsMale * Age + Rank * YAR$^2$ + Ret*** | **16** | **3958.7** | **4072.3** |
| *Attrition* | *IsMale * (Age + Rank) + Rank * YAR$^2$ + Ret* | 19 | 3959.1 | 4094.0 |
| *Attrition* | *IsMale * (Age + Rank * YAR$^2$ + Ret + IsSTEM)* | 30 | 3963.4 | 4176.3 |
| *Attrition* | *Rank * YAR$^2$ + Ret + IsSTEM* | 14 | 3966.0 | 4065.4 |
| *Attrition* | *IsMale * (Age + Rank) + Rank * YAR$^2$ + Ret* | 13 | 3970.6 | 4062.9 |
| *Promotion* | *IsMale * (Age + Rank) + Rank * YAR$^2$ + Ret* | 15 | 3973.9 | 4077.5 |
| *Promotion* | *Rank * YAR$^2$ + Ret + IsSTEM* | 11 | 4012.4 | 4088.5 |
| *Attrition* | *IsMale * (Age + Rank)* | 10 | 4042.0 | 4113.0 |
| *Attrition* | *IsMale * (Age + Rank) + IsSTEM* | 11 | 4042.7 | 4120.8 |
| *Promotion* | *Rank * YAR$^2$ + Ret* | 10 | 4132.3 | 4201.4 |
| *Promotion* | *IsMale * (Age + Rank) + IsSTEM* | 9 | 4208.7 | 4271.0 |
| *Attrition* | *IsSTEM* | 2 | 4209.9 | 4224.1 |
| *Promotion* | *IsMale * (Age + Rank)* | 8 | 4267.3 | 4322.6 |
| *Promotion* | *IsSTEM* | 2 | 4316.2 | 4330.0 |

Table S3: **Generalised linear regression models of promotion and attrition fit using University of Canterbury data and sorted by ascending AIC score**. An AIC difference of $\Delta\mathrm{AIC} = 2$ was considered significant. Numbers of parameters $k$ includes a baseline term. *YearsAtRank* is abbreviated to *YAR* and *Retirement* is abbreviated to *Ret*.

### 4. References


1. Cech, E.A. and T.J. Waidzunas, *Systemic inequalities for LGBTQ professionals in STEM.* Sci Adv, 2021. **7**(3).
2. Moss-Racusin, C.A., et al., *Boosting the Sustainable Representation of Women in STEM With Evidence-Based Policy Initiatives.* Policy Insights from the Behavioral and Brain Sciences, 2021.
3. Moss-Racusin, C.A., et al., *Gender Bias Produces Gender Gaps in STEM Engagement.* Sex Roles, 2018.
4. Thomas, N.R., D.J. Poole, and J.M. Herbers, *Gender in Science and Engineering Faculties: Demographic Inertia Revisited.* PLoS One, 2015. **10**(10): p. e0139767.
5. Canterbury, U.o. *General Staff Collective Employment Agreement*. 2024 [cited 2026 04/03/2026].
6. Moss-Racusin, C.A., et al., *Science faculty's subtle gender biases favor male students.* Proc Natl Acad Sci U S A, 2012. **109**(41): p. 16474-9.
7. Mayer, S.J. and J.M.K. Rathmann, *How does research productivity relate to gender? Analyzing gender differences for multiple publication dimensions.* Scientometrics, 2018. **117**(3): p. 1663-1693.
8. Nygaard, L.P., D.W. Aksnes, and F.N. Piro, *Identifying gender disparities in research performance: the importance of comparing apples with apples.* Higher Education, 2022. **84**(5): p. 1127-1142.
9. van den Besselaar, P. and U. Sandstrom, *Vicious circles of gender bias, lower positions, and lower performance: Gender differences in scholarly productivity and impact.* PLoS One, 2017. **12**(8): p. e0183301.
10. Abramo, G., C.A. D'Angelo, and A. Caprasecca, *Gender differences in research productivity: A bibliometric analysis of the Italian academic system.* Scientometrics, 2009. **79**(3): p. 517-539.
11. Benjamin, E., *Disparities in the Salaries and Appointments of Academic Women and Men.* Academe, 1999. **85**(1): p. 60-62.
12. Costas, R., T.N. van Leeuwen, and M. Bordons, *A bibliometric classificatory approach for the study and assessment of research performance at the individual level: The effects of age on productivity and impact.* Journal of the American Society for Information Science and Technology, 2010. **61**(8): p. 1564-1581.
13. Huang, J., et al., *Historical comparison of gender inequality in scientific careers across countries and disciplines.* Proc Natl Acad Sci U S A, 2020. **117**(9): p. 4609-4616.
14. Manfredi, S.L.a.S., *Balancing Gender in Higher Education.* The European Journal ofWomen's Studies, 2000. **7**(33).
15. Brower, A. and A. James, *Research performance and age explain less than half of the gender pay gap in New Zealand universities.* PLoS ONE, 2020. **15**(1): p. 1--13.
